# Multi-omics analysis of long-term cultured human islets

**DOI:** 10.1101/2024.12.25.626491

**Authors:** Guihua Sun, Meirigeng Qi, Olivia Sun, Elizabeth M. Lizhar, Deborah Hussey, Yanhong Shi, Arthur D. Riggs

## Abstract

β-cell dysfunction in pancreatic islets, characterized as either the loss of β-cell mass or the resistance of β-cell to glucose, is the leading cause of progression to diabetes. Islet transplantation became a promising approach to replenish functional β-cell mass. However, not much known about changes in islets used for transplantation after isolation. We have subjected human islets into long-term *in vitro* culture (LTC) and characterized those survived islets. While most of the dysregulated genes were downregulated during LTC, specific groups of mRNA or miRNA were upregulated, and they are involved in specific pathways. In general, α-cells and β-cells of LTC-islets have elevated expressions of *MAFB* and *MAFA* genes, respectively. We also found that exocrine cells were eliminated faster than endocrine cells, and β-cells were lost at a higher rate than α-cells. Interestingly, one specific group of cells that have characteristics of immature α-cells or β-cells, were enriched in LTC-islets, revealing the possibility of transdifferentiation of α-cells to β-cells, or dedifferentiation of β-cells to α -cells, under *in vitro* culture. Our results suggest that there are intrinsic cellular and molecular mechanisms in pancreatic cells that are associated with their maturity and correlated with their survival ability under unfavorable living conditions.

## Introduction

Diabetes mellitus and its associated complications affect billions of people around the world, and the heavy health and economic burden to society caused by this disease is on the rise. Diabetes can be classified into two major classes based on the mechanism of how they lose their functional β-cells, which is a specific type of cell in the pancreatic islets that can produce insulin for the metabolism of glucose. Type I diabetes (T_1_D) is resulted from a deficiency in β-cell mass because most of their β-cells were believed to be destroyed by autoimmune reactions. In Type 2 diabetes (T_2_D), β-cells lose their effective responses to glucose stimulus and are resisted to insulin signaling ^1^. Despite significant advances in diabetes research and therapies for diabetes, an ideal cure remains elusive, likely because the mechanisms behind β-cell death in T_1_D and β-cell damage in T_2_D are not yet fully understood. Islet transplantation became a promising approach to treat patients with T1D because it provides optimal glycemic control, higher survival rate, and better quality of life for patients ^1–4^. However, in addition to the costly and invasive procedure, patients will face the consequence of toxic immune reactions and other restraints.

Moreover, the major hurdle is that there is only a limited number of human pancreases available from cardiac donors. Islets of Langerhans are mini-organs, cryopreserved islets or islets that have been cultured for a longer period, are usually not suitable for islet transplantation ^2,4–8^. Although a great progress has been made in the cryopreservation of human islets ^8^, *in vitro* culture of human islets, which will subject islets to stress, is still a necessary and critical step in islet transplantation, so that the quality and quantity of donor islets can be evaluated, and it also provides additional time to get recipients ready for the operation. *In vitro* culture of islets will also allow researchers to study their function, to characterize the subtype of cells in human islets, and to study the function of pancreatic genes or subtypes of cells. While islets transplantation only needs islets to be cultured *in vitro* for less than 72 hours, islets used for research may need to be in culture for a longer time to define the functions and features of human islets and subtypes of cells in islets. Genetic engineering to obtain islets with enhanced or gained function may also require the islets to be cultured for an extended time. Therefore, there is a need for novel protocols of long-term *in vitro* culture (LTC) of islets that can preserve their function. We hypothesized that systemic genes profiling and subtype cells characterization of LTC-islets may yield knowledge of how cells in islets can survive LTC to aid the development of novel islet culture protocols. To investigate problems related to LTC-islets as well as mechanisms underlaying β-cell death or loss of function at both molecular and cellular levels, we have assessed gene profiling changes using bulk RNA sequencing (RNA-seq), small RNA changes using small RNA sequencing (smRNA-seq), and cell composition changes using single cell RNA sequencing (scRNA-seq) for LTC-islets. We also compared the surviving LTC-islets with islets from T_1_D and T_2_D patients. Our results showed that both subtype cells and gene profiling underwent dramatic changes in islets after LTC. While LTC-islets in some respects resembled more islets of T_2_D than islets of T_1_D or freshly isolated islets, cells in LTC-islets exhibited different gene profiling from islets of T_1_D or T_2_D in that LTC-islets showed higher expression of specific genes or miRNAs. We observed that exocrines cells were eliminated at a higher rate than endocrine cells which may have a major impact on the function of LTC-islets. We also observed that specific groups of α-cells and β-like-cells have elevated expression of *MAFB* and *MAFA* genes, respectively, under LTC. Our data demonstrated that α-cells have a better chance to survive LTC than β-cells, supporting an idea that β-cells have specific intrinsic features that make them less resilient to unfavorable survival conditions, which is independent of β-cell death caused by autoimmune attacks occurring in T_1_D. Our study has provided an alternative molecular and cellular basis of β-cell destruction and might aid the development of novel procedures of generating functional LTC-islets that will benefit both islet transplantation and research.

## Results

### Some human islet cells can survive long term *in vitro* culture

*In vitro* cultured human islets are used in both islet transplantation and research experiments, and progressions in islet transplantation are critical for T_1_D therapy ^3,5^. Previous work showed that isolated adult rat islets of Langerhans can survive 97 days in an artificial capillary culture unit without loss of functionality ^9,10^. On the other hand, human islets failed to preserve their function and were not suitable for islets transplantation after being cultured *in vitro* for a long period ^2,5–7^. The reason behind the difference between human islets and rat islets remains elusive. There is limited cellular and molecular data on the changes that occur in human islets during *in vitro* culture, as well as on how long they can be cultured without significant loss of normal function. In the current study, we first examined islet survival during *in vitro* culture. To track the survival and proliferation of α and β-cells in LTC, human islets were dissociated and transduced by lentiviral vectors expressing EGFP driven by the human insulin (INS) promoter and tdTomato driven by the CMV promoter (INS-EGFP-CMV-tdTomato) to track β-cells, or mCherry driven by the human glucagon (GCG) promoter and EGFP driven by the CMV promoter (GCG-tdTomato-CMV-EGFP) to track α-cells. We observed that the cell viability is high (90% to 98.7%) after islet dissociation, most cells aggregated and survived at week-2, about 50% survived at week-3, but over 70% of cell death happened around week-4, and less than 5% were left after 46 days in LTC (**Fig. 1A**, **Table S1**). These results suggest that two weeks may be the time limit for human islets to maintain their cell population under LTC. We decided to study gene profiling changes and subtype cells changes in human islets cultured up to four weeks using both scRNA-seq and bulk RNA-seq to characterize cells in LTC-islets. Although islets are typically cultured at 37 °C and/or 22 °C before transplantation ^6^, we only tested islets cultured at 37 °C to assess long-term effects as islets are unlikely to remain viable for extended periods at 22 °C. To eliminate the potential confounding effects from different accessibility to culture medium between cells outside islets and cells inside islets, dissociated human islets were used for all studies.

**Figure 1.**
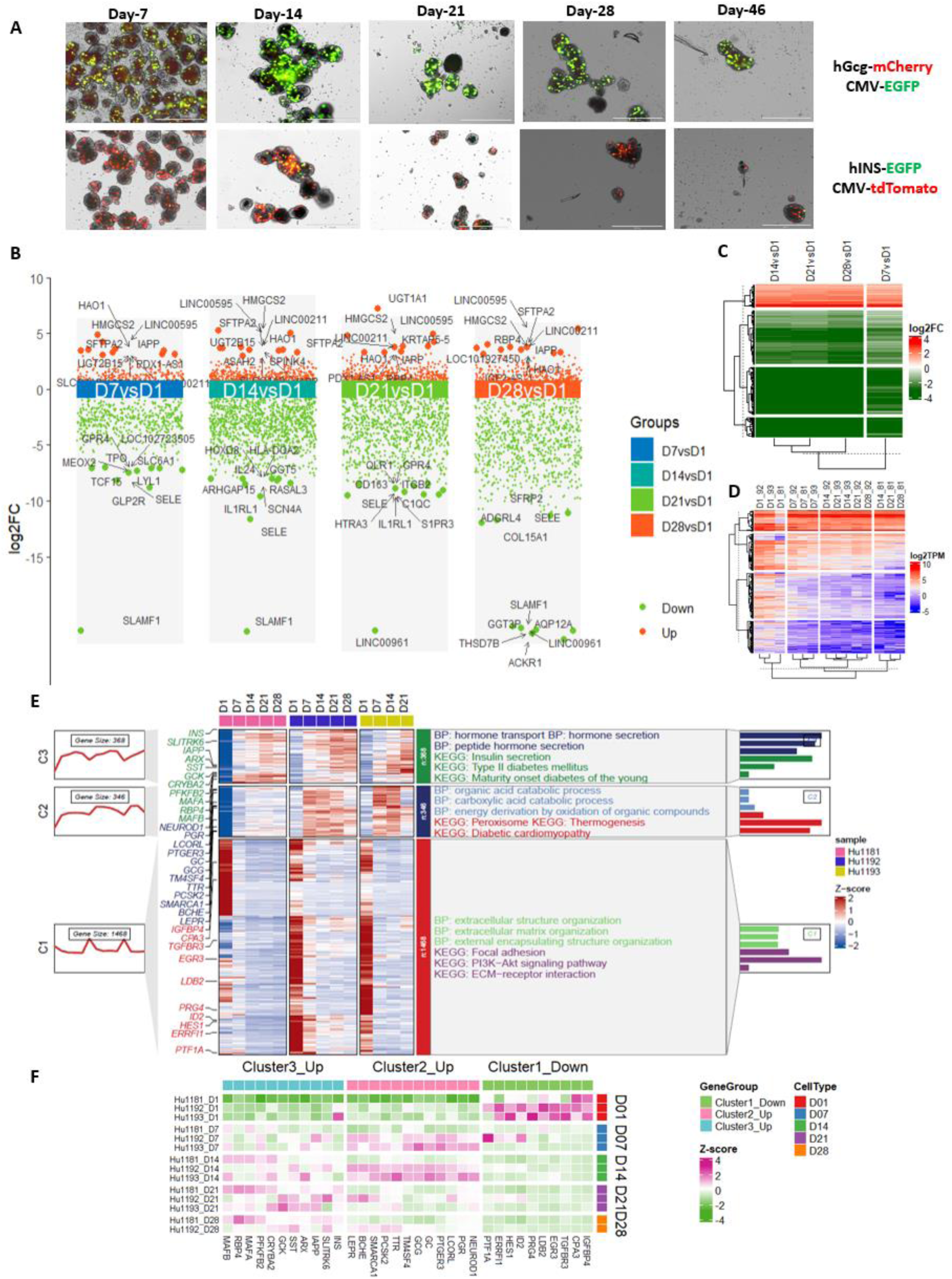
RNA-seq analyses of LTC islets. **A.** Tracking α and β-cells in long-term *in vitro* cultured human islets by reporters. A lentiviral vector with human glucagon (hGCG) promoter drive mCherry and CMV promoter drive EGFP (hGCG-mCherry-CMV-EGFP) was used to track α-cells. A lentiviral vector with human insulin (hINS) promoter drive EGFP and CMV promoter drive tdTomato (hINS-EGFP-CMV-tdTomato) was used to track β-cells. **B.** A comparison plot of DEGs between D7, D14, D21, or D28 and D1 (D7vsD1, D14vsD1, D14vsD1, and D28vsD1). Top-10 upregulated genes and top-10 downregulated genes were labeled by their names and marked by larger red or green dots respectively. **C.** A heatmap plot showed clusters of log2 fold changes of genes between D7, D14, D21, or D28 and D1. **D.** A heatmap plot of gene expression measured by log2 TPM in all samples. **E.** A combinatorial plot of DEGs between D7, D14, D21, or D28 and D1 in all samples. The heatmap showed normalized gene expressions in all samples. DEGs that are also pancreatic essential genes were labeled on the left side of the heatmap. Clusters of DEGs and their associated bioprocesses (GO:BP) and KEGG pathways were plotted on the right side of the heatmap. The trend of gene expression in each cluster were plotted on the far-left side as line graphs and the gene sets size associated with each BP and KEGG were plotted on the far-right side as bar graphs. **F.** A dedicated heatmap plot of DEGs that are also pancreatic essential genes in E. Normalized gene expression data were used for this plot.

### Bulk RNA-seq revealed gene profiling changes in LTC-islets

To observe gene expression changes in LTC-islets, we first performed bulk RNA-seq of LTC-islets from individual donors at different time points of culture. The islet cells from three donors were cultured for up to four weeks. Total RNAs were collected weekly. RNA-seq results from samples at each culture time point (D7, D14, D21, and D28 for samples at day 7, day 14, day 21, and day 28, respectively) were compared to samples that were collected shortly after islet isolation and cultured for only a few hours (samples of day-1, hereafter referred to as D1, representing islets prepared for transplantation) to identify differentially expressed genes (DEGs) at each time point. DEGs with significant changes between each time point to D1 were used to identify significantly and consistently changed genes during LTC.

We observed that more genes were significantly downregulated than genes that were upregulated which corresponds to the death of many cells during LTC. Among the top-10 upregulated genes, *IAPP*, *HAO1*, *HMGCS2*, *SFTPA2*, *LINC00595*, and *LINC00211* are present in all four time points. However, among the top-10 downregulated genes, only *SELE* is present in all four time points and *SLAMF1* is present in three time points (**Fig. 1B**, **S1**). Therefore, among the top-10 genes, more were consistently upregulated than downregulated across all four time points.

A subgroup of genes from all DEGs in all four time points were used to cluster DEGs by fold change and transcript abundance was measured by transcripts per million reads (TPM). Based on the clustering result from fold change data, D14 vs D1, D21 vs D1, and D28 vs D1 were clustered together, and they were separated from D7 vs D1 (**Fig. 1C**). Based on the clustering result from TPM of genes, samples of D1 were in a cluster, samples of D7 were in a cluster, and samples of D14, D21, and D28 were classified into another two clusters (**Fig. 1D**). Both fold-change data and gene expression heatmap agreed with DEG analysis data that there are more downregulated genes than upregulated genes (**Fig. 1C-D**). The above results indicated that islets could go through significant gene expression changes in the first two weeks of LTC and LTC-islets tend to have similar gene expression profiles after two weeks and beyond (**Fig. 1B-D**).

We checked the expression of 129 essential pancreatic genes (defined as genes which encoding pancreatic hormones or transcription factors) in the pooled DEGs (**Supplementary file 1**) and found 32 of them were present (*INS* was added for comparison). Therefore, a large percentage of essential genes in the pancreas were affected by LTC. Cluster analysis showed that these essential pancreatic genes fit into three clusters. Clusters 2 and 3 contain 22 upregulated genes and cluster 1 contains 10 downregulated genes. Here again there were more upregulated than downregulated essential pancreatic genes in the DEGs which contrasted with the result that more genes were downregulated globally (**Fig. 1B-F**, **S1**). Gene ontology (GO) and KEGG pathway analysis showed that hormone secretion, carboxylic acid metabolism, extracellular structure and matrix organization, focal adhesion, PI3K-AKT signaling pathway, and ECM-receptors are among those top affected bioprocesses and pathways (**Fig. 1E**).

Next, we performed a more detailed pathway analysis using DEGs from the four time points. Based on the analysis results of DEGs of D7 vs D1, we found that many DEGs are involved in bioprocesses of the extracellular matrix and structure, cell adhesion, and molecular activities of several ion channels, indicating that *in vitro* culture will affect islets on cell organization and cellular activities (**Fig. 2A-C**). Comparison pathway analysis using all DEGs at each time point showed all 13 major altered pathways at D7 were also present in other time points. Additional pathways were affected in and after the second week of culture and most of these pathways remained as affected pathways at later time points. Importantly, pathways related to immune activities, such as TNF signaling, cytokines and cytokine receptor interactions, showed up after the D7 time point (**Fig. 2D**). This pathway analysis revealed that, under culture conditions, genes involved in interactions between islets cells were affected and altered genes were involved in the cell cycle, cell metabolism, growth, proliferation, and survival, and most likely immune activities were triggered before D7. Therefore, we can conclude that under LTC, gene expression in islets will go through many changes in the first week, but major changes will be happened in the second week, the dysregulated genes are involved in many pathways, and immune responses will be triggered within a week in culture. Next, we performed a comparison pathway analysis using only upregulated DEGs at each time point. These genes are of particular interest because they were probably activated to mediate cell survival during LTC. The top two pathways are insulin and bile secretions, and they are present across all time points. However, we observed N-glycan biogenesis, carbohydrate digestion and absorption, and the cAMP signaling pathway were largely affected in D7 and D14, but not D21 or D28. These pathways were probably specifically activated during the first two weeks of islets that are first grown *in vitro* and were shut off thereafter, hence genes in these pathways may mediate islets’ survival in the early stages of LTC (**Fig. 2E**).

**Figure 2.**
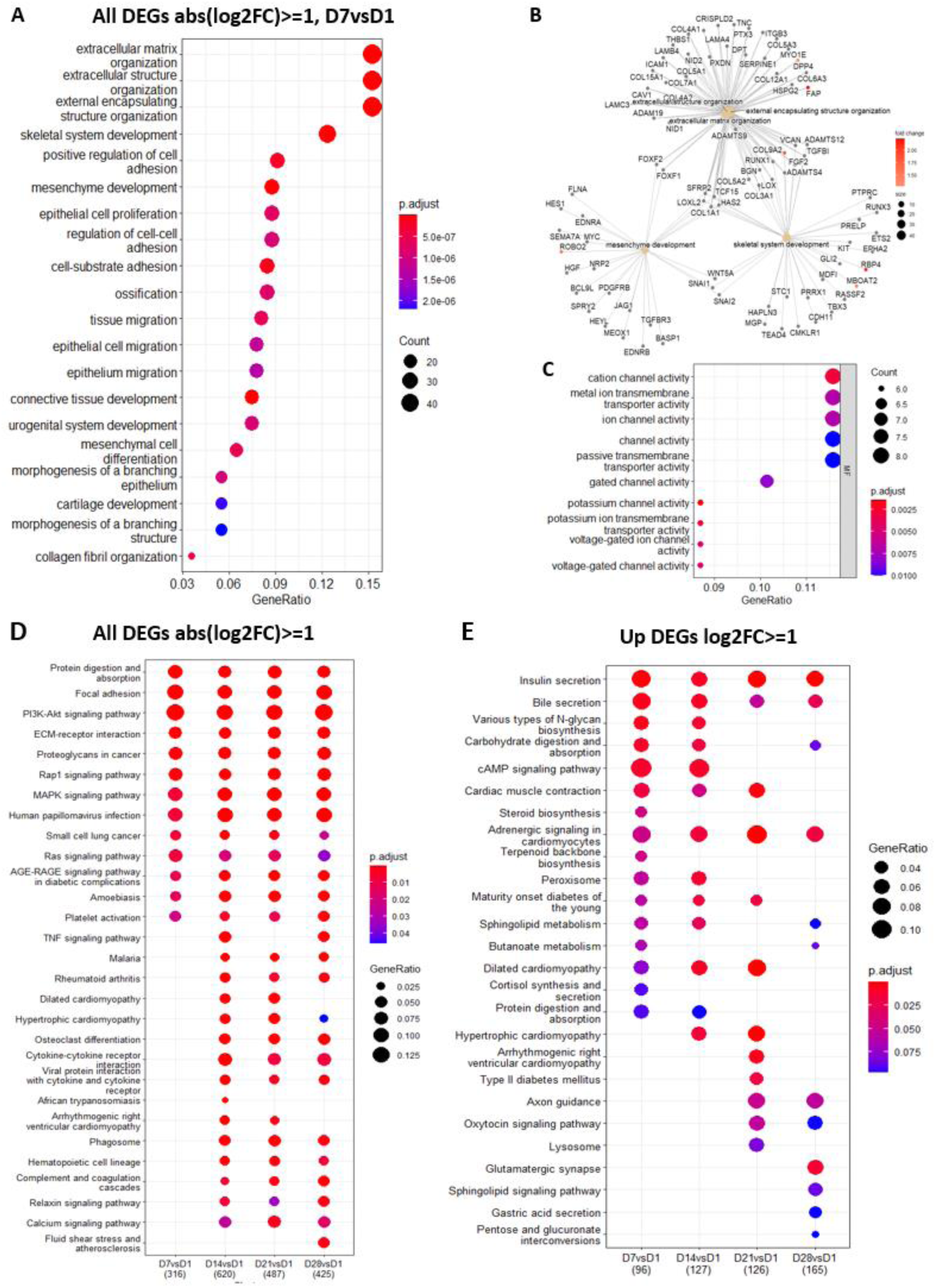
GO bioprocess analyses of RNA-seq of LTC islets. **A.** A dot plot showed top-25 GO-BPs using DEGs (|log2FC|>=1) of D7vsD1. **B.** A network plot of top-5 categories of GO-BPs in A. along with their associated DEGs. **C.** A dot plot of top-10 GO molecular functions (GO:MF) involved by DEGs in A. **D.** A dot plot showed the comparison pathway analysis of involvements of DEGs from D7vsD1, D14vsD1, D21vsD1, and D28vsD1. **E.** A dot plot showed the comparison pathway analysis of involvements of upregulated DEGs from D7vsD1, D14vsD1, D21vsD1, and D28vsD1.

Among all upregulated DEGs, *IAPP, ARX, SST, GCK, MAFA,* and *MAFB* genes are in the clusters of upregulated pancreatic essential genes and were upregulated consistently at all time points (**Fig. 1E-F**). We paid special attention to the *IAPP* gene, which encodes amylin, a hormone mainly expressed in β-cells and co-secreted with insulin in the ratio of approximately 100:1 (insulin:amylin). Published data suggest amylin and insulin are regulated by similar factors ^11^. However, *INS* gene expression was slightly upregulated in the first two weeks of LTC then mildly upregulated during the last two weeks of LTC while expressions of the *IAPP* gene were constantly significantly upregulated (**Fig. 1&S1**), indicating that β-cells produce excess amylin, and the normal ratio of insulin to amylin was disrupted under LTC. Upregulated expression of *IAPP* may be one of the major reasons for the death of β-cells in LTC-islets because published studies suggest that excess amylin can trigger destruction to β-cells ^12,13^.

### smRNA-seq revealed miRNA changes in LTC-islets

Beside transcription factors, pancreas development and cell differentiation are also controlled by miRNAs, a family of small non-coding RNAs with various regulatory roles in many biological processes^14^. We performed smRNA-seq of the same samples that were used for the above RNA-seq. Like gene expression changes in the RNA-seq data, significantly downregulated differentially expressed miRNAs (DEmiRs) also exceeded the number of significantly upregulated DEmiRs in about a ratio of 6 to 1 (64 versus 10) (**Fig. 3A**, **Supplementary file 1**). Based on the clustering data using fold-change of DEmiRs, DEmiRs separated islets that were cultured for a week (D7 vs D1) from the rest of the LTC-islets (D14 vs D1, D21 vs D1, and D28 vs D1). Clustering miRNAs using read count per million reads (RPM) measured miRNA abundance, samples of D1 were clustered together, samples of D7 were closely clustered, and the rest of the samples (D14, D21, and D28) were closely clustered (**Fig. 3B-C**). Therefore, despite the much smaller population of miRNAs compared to mRNAs, the miRNA profiling results also indicated islets can go through major miRNA gene expression changes within the first week of culturing, and islets exhibited similar miRNA profiles in later time points after two weeks in LTC.

**Figure 3.**
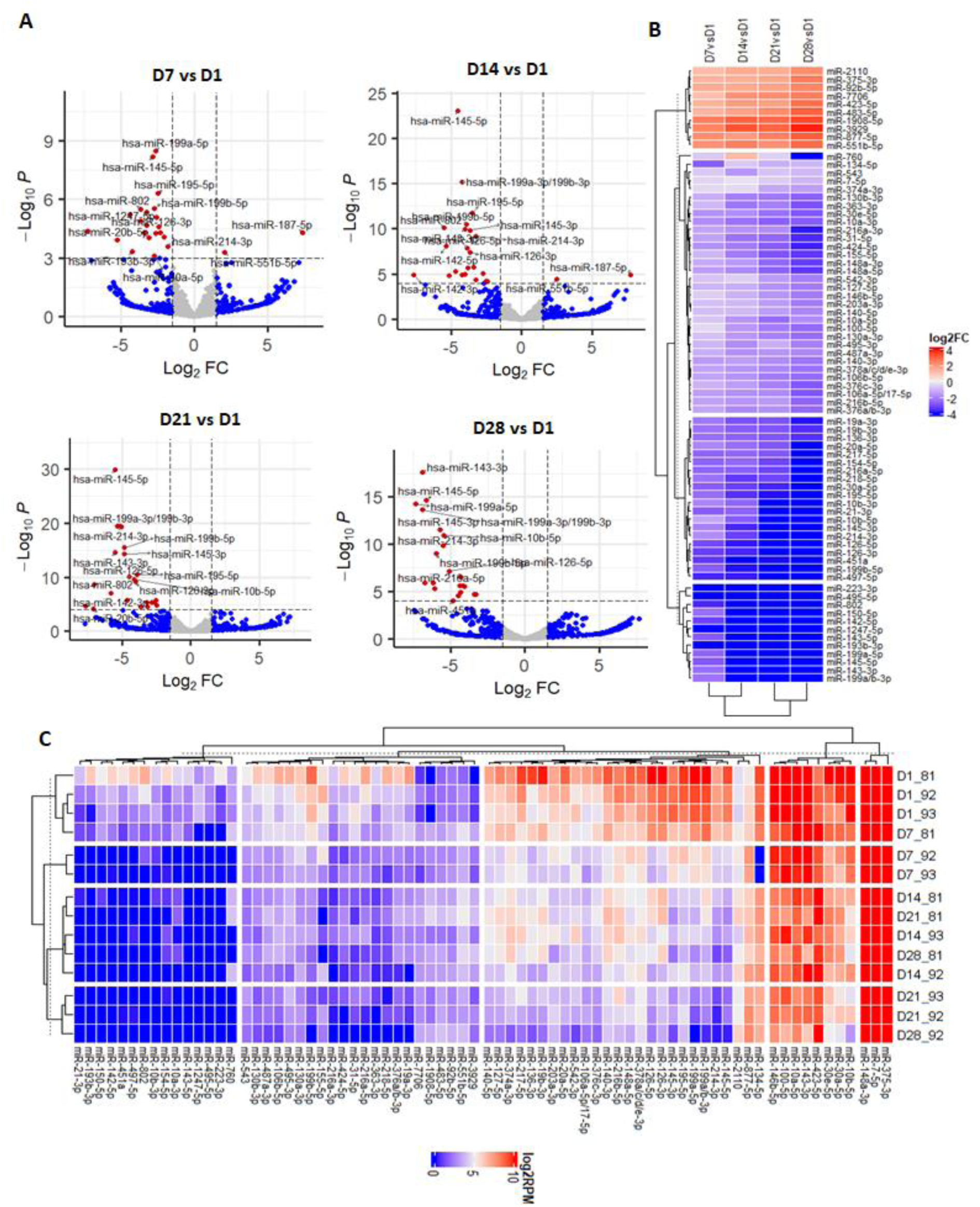
smRNA-seq analyses of LTC-islets. **A.** Volcano plots of differential expressed miRNAs (DEmiRs) of D7vsD1, D14vsD1, D14vsD1, and D28vsD1. **B.** A heatmap plot showed clusters of log2 fold changes of DEmiRs between D7, D14, D21, or D28 and D1. **C.** A heatmap plot showed clusters of DEmiRs between D7, D14, D21, or D28 and D1. Log2 transformed read per million miRNA reads (RPM) were used for this plot.

Among the three most abundant human pancreas miRNAs, miRNA-375-3p, miR-7-5p and miR-148a-3p, the abundance of miRNA-375-3p increased with the culture duration. In contrast, the abundances of miR-7-5p and miR-148a-3p decreased, although only the decrease in miR-148a-3p was statistically significant (*p-adj* < 0.05) (**Fig. 3B**, **Supplementary file 1**). The decrease of miR-148a-3p is probably caused by the loss of acinar cells because miR-148a-3p is the most abundant miRNA in the human acinar cells ^15^. According to published miRNA profiling data in sorted α and β-cells (GSE52314) ^15,16^, miR-375-3p is expressed at a higher level in α-cells than in β-cells (43.47% versus 31.83% of all miRNAs), hence the increase of miR-375-3p, which coincides with the increase of *ARX, GCG, MAFB* genes, marker genes of α-cells, in RNA-seq data (**Fig. 1F**). Among the significantly upregulated DEmiRs, only miR-877-5p and miR-423-5p were expressed at high levels and they also had a similar increasing pattern as miR-375-3p in that their expression levels were increased with culture time (**Fig. 3B-C**). Among the downregulated DEmiRs, more than a dozen of them (miR-10a/b-5p, miR-19b-3p, miR-30a/e-5p, miR-100-5p, miR-126-5p/3p, miR-136-3p, miR-140-3p, miR-143-3p, miR-145-5/3p, miR-146b-5p, miR-148a-5/3p, miR-195-5p, miR-199a/b-3p/5p, miR-216b-5p, miR-217-5p, miR-378a/c/d/e-3p) have high expression levels and their expression levels were also decreased with culture time. The downregulation of miR-146b-5p, an immune-response-related miRNA, suggests that LTC-islets may lose some of their immune response activities during culture.

We performed a GO and KEGG pathway enrichment analysis for the above dysregulated miRNAs. We first focused on the two upregulated DEmiRs, miR-877-5p and miR-423-5p, which may be the two important miRNAs for supporting islets survival during LTC. Results showed that they are mainly involved in antifungal related bioprocesses (**Fig. S2**). Therefore, the antifungal innate immune responses were activated during the full culture period which was not specifically revealed by RNA-seq results. Next, we performed a similar analysis for those downregulated DEmiRs. We found the cell proliferation, cell adhesion, and PI3K-AKT signaling pathways were affected. This observation aligned with the effects from upregulated DEGs, and the primary function of miRNAs, which is to downregulate target genes (**Fig. S3A-B**). Among all the protein-protein interactions formed by targets of downregulated DEmiRs, one that is involved two important pancreatic genes, *IRS1* and *IGF1R*, stood out (**Fig. S3C**). According to TargetScan 8.0, *IRS1* is a conserved target among vertebrates for miR-7-5p, miR-10-5p, miR-126-3p, and miR-145-5p, while *IGF1R* is a conserved target among vertebrates for miR-7-5p, miR-30-5p, miR-100-5p, miR-143-3p, and miR-145-5p ^17^ (**Fig. S3**). Here the DEmiRs showed their advantages over DEGs as DEmiRs revealed LTC-islets were under threat from fungal infection and pointed out pathways involving insulin, IRS1, and IGF1R were activated during LTC.

### LTC-islets are different from residual islets of T_1_D

Given the significant cell loss in LTC-islets, which resembles the cell death-mediated cell loss observed in T_1_D, we investigated whether LTC-islets are similar to residual islets in T_1_D. To address this, we compared mRNA and miRNA profiling data between LTC-islets and T_1_D islets.

We first downloaded an RNA-seq dataset of T_1_D (GSE162689 from GEO database) and performed DEG analysis using this dataset ^18^. DEGs of LTC-islets and T_1_D-islets showed a clear difference that the two-upregulated genes in LTC-islets, *MAFA* and *IAPP,* were among the significantly downregulated genes in T_1_D-islets (**Fig**. **S4**). While *INS* was downregulated in both T_1_D-islets and LTC-islets, *INS* was downregulated at a much larger scale in T_1_D-islets than in LTC-islets (**Fig. 1F**, **S4**). Next, to compare DEmiRs between LTC-islets and T_1_D-islets, we used a published miRNA profiling result from tissues of T_1_D ^19,20^. Among all highly expressed miRNAs in islets, only miR-375-3p was upregulated in both LTC-islets and T_1_D-islets. In contrast, miR-10a-5p and miR-148a-3p were upregulated in T_1_D-islets but downregulated in LTC-islets, while miR-100-5p and miR-146a-5p were downregulated in both T_1_D-islets and LTC-islets ^19,20^. Hence, DEmiRs exhibited differences in LTC-islets and T_1_D-islets, and islets lost protection from miR-146a-5p in both T_1_D-islets and LTC-islets.

In summary, both DEGs and DEmiRs indicated that LTC-islets have a major difference from residual islets in T_1_D. The plot of essential pancreatic genes and pathway analysis suggest that LTC-islets may be more closely resemble islets of T_2_D and maturity onset Diabetes of the young (**Fig. 1E**).

### Cell composition of LTC-islets differed from islets of T_1_D or T_2_D

The differences observed in RNA-seq and smRNA-seq results between LTC-islets and diabetes patients’ samples prompted us to conduct a more detailed characterization of cells in LTC-islets. This further analysis aimed to deeply examine how these cells differ from the residual islets found in T_1_D and T_2_D. We performed scRNA-seq to compare the cell compositions between islets that have been cultured for a short-term and long-term. Human islets were cultured for less than one day after isolation before being dissociated into single cells. An aliquot of dissociated cells was used for scRNA-seq on the same day right after dissociation (hereafter we will refer to this sample as Day1). Remaining cells were cultured for four weeks. On the final day of LTC, residual cells were dissociated and sequenced (hereafter we will refer to this sample as Day29).

We first analyzed the scRNA-seq using too-many-cells, a scRNA-seq data analysis tool that will produce a tree like structure of all clusters of cells for a nice and easy visual comparison of all cells and their relationships ^21^. There are a total of 884 clusters or nodes and some subtrees/nodes only present either in samples of Day1 or Day 29, indicating that some groups of cells were lost, and some groups of cells were raised or transformed during LTC (**Fig. 4A**). Then we trimmed (smart-cutoff 2, min-size 1) the tree into a smaller tree (58 nodes) that is easy to work with and has better visibility (**Fig. 4B-C**). We identified the cell types associated with each branch by plotting of the expression of marker genes (**Fig. 4B-P**). Cells of parental node 40 (PN40) were *CPA1*-positive cells and were mainly present in the Day1 sample, indicating that there is a major loss of Acinar cells in LTC. Similarly, PN20 cells are *KRT19*-positive and mainly present in the Day1 sample, indicating that there is also a major loss of ductal cells. This result is unexpected because exocrine cells are generally considered more resilient than endocrine cells under unfavorable conditions, and it was anticipated that more of them would survive LTC. Cells in the PN27 branch have a similar cell distribution in Day1 and Day29 samples, and lacked pancreatic marker gene expression, suggesting that they are probably non-pancreatic cells or transformed cells that lost the expression of pancreatic marker genes. Cells of PN3 are probably endocrine cells. Plots of marker genes of β-cells (*INS*/*IAPP*/*MAFA*/*PAX6*) and *GCG* overlaid plot with *INS* indicated Node 19 (N19) and N13 were both *INS*/*MAFA-*positive cells with relatively low *GCG* expression, indicating that they are mainly β-cells (**Fig. 4C, D, F, H, P, J**). Plots of marker genes of α-cells (*GCG*/*ARX*/*MAFB*/*PAX6*) indicated that cells of PN4 are mainly α-cells (**Fig. 4C, E, G, I, P**). While both cells of PN5 and PN12, derivatives of PN4, were *GCG*/*ARX*-positive, cells of PN12 were mainly present in sample of Day1 and cells of PN5 were mainly present in sample of Day29. These results indicating that, although these cells are close to each other, they exhibit major changes during LTC and that cells of PN5 were at an earlier stage of development than PN12. One derivative branch of PN12, N13, represented an interesting group of cells that were mainly present in the Day1 sample, and showed higher expression of INS and lower expression of GCG (**Fig. 4J**). These cells could be classified as β-cells, but they are among the branches of α-cells and were lost during LTC because they mainly present in the Day1 sample. Cells of PN5, mainly in the Day29 sample, also have a higher expression of *MAFB* compared to cells of PN12, indicating that some α-cells could gain higher *MAFB* expression during LTC. On the contrary, cells in P19 had a higher expression of *MAFA* and *IAPP* than cells in P13, indicating that some β-cells could gain higher *MAFA* expression during LTC. These results demonstrate that LTC induces major changes in both the cell composition and gene expression in islets.

**Figure 4.**
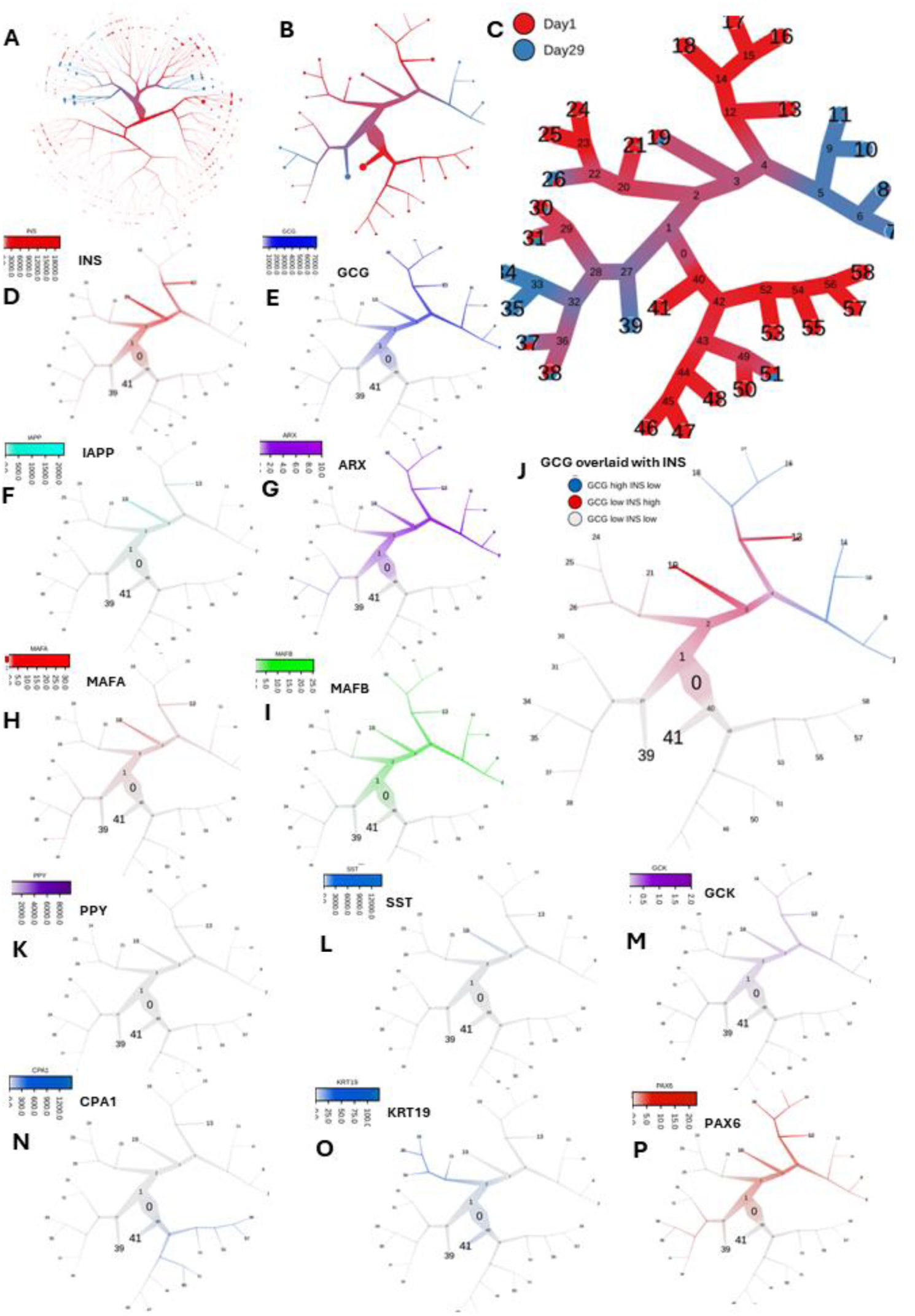
scRNA-seq analyses by too-many-cells. **A.** too-many-cells predicted cells distribution tree of Day1 and Day29 samples. **B & C**. Trimmed tree. **D to I & K to P**: Plots of cell marker genes on the trimmed tree. **J**. Overlap plot of INS with GCG on the trimmed tree.

Next, we performed a more detailed analysis of our datasets using Seurat and compared them to scRNA-seq datasets of T_1_D and T_2_D. We performed integrated scRNA-seq data analysis on our scRNA-seq datasets with published scRNA-seq datasets of T_1_D (GSE148073) ^22^ and T_2_D (GSE816085) ^23^ patient islets using the integrated analysis workflow described in Seurat ^24^ (**Fig. S5**). Using the annotated cell types in scRNA-seq datasets of human pancreas (panc8) that were curated in SeuratData as a reference ^24^, we predicted cells in all four datasets (Day1, Day29, T_1_D, and T_2_D) and the prediction was successful as indicated by the expression of marker genes in the predicted α-cells (*GCG, ARX, MAFB)* and β-cells (*INS, IAPP, MAFA*) (**Fig. 5A-B**). There were 22 predicted clusters with unique gene expression signatures and cells were annotated as 13 subtypes of cells. Besides the expected large population of pancreatic cells, there were also non-pancreas specific cells such as stellate, endothelial, macrophage, mast, and Schwann cells (**Fig. 5C-D**).

**Figure 5.**
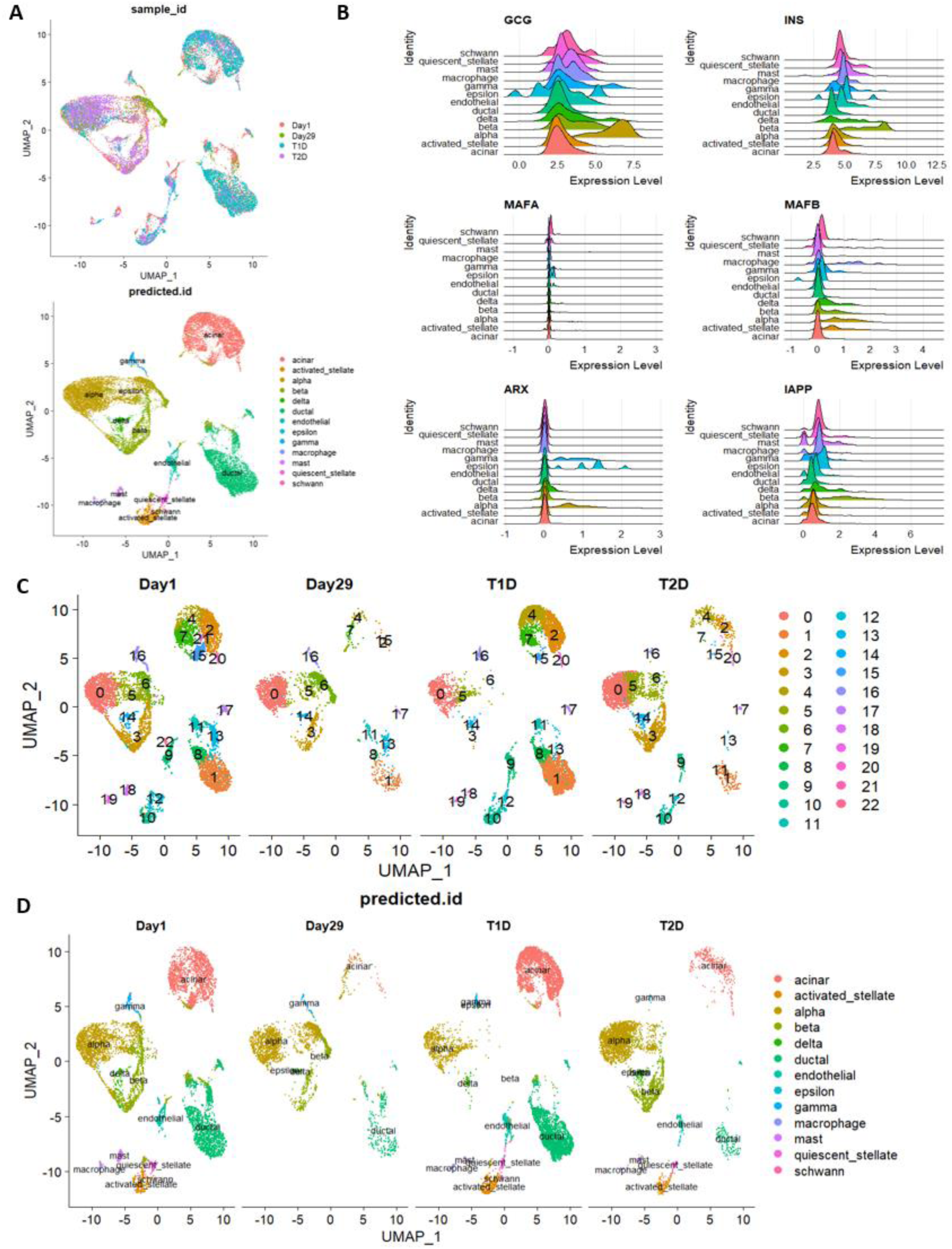
scRNA-seq analyses of LTC-islets—predicted cell clusters and cell types-Seurat. **A.** Upper panel: an overplay plot to show cell distributions in each group of samples and cells were colored by names of the group; Lower panel: a cell distribution plot to show cell type distributions in all samples and cells were colored by their predicted cell types. **B.** Ridge plots of the expression ranges of some marker genes of α-cells (*GCG, ARX, MAFB*) and β-cells (*INS, IAPP, MAFA)* in all predicted cell types. **C.** Side-to-side comparison plots of cells with their predicated clusters in each group of samples and cells were colored by their predicated clusters. **D.** Side-to-side comparison plots of cells with their predicated cell type in each group of samples and cells were colored by their predicated cell types.

The side-by-side comparison plotting by samples clearly showed different cell composition among all four sample groups according to plots by predicted cell clusters and cell types (**Fig. 5C-D**). The plots of expression ranges of marker genes in six major pancreatic cell types among all predicted clusters showed heterogeneity in α-cells, β-cells, acinar cells (*CPA1-* positive cells), and ductal cells (*KRT19*-positive cells), agreeing with previous reports (**Fig. 5A**) ^25,26^. Next, we performed a more detailed analysis to dissect the correlations among cell types, cell clusters, and marker genes.

### Cells of Day29 lost the majority of exocrine cells

Comparing sample of Day29 to sample of Day1, we observed that, in addition to the expected large population of endocrine cells, a substantial number of exocrine cells were present in these islets despite their estimated purity scores being over 80%, as measured by a traditional islets staining approach (**Table S1**) ^27^. In agreement with the too-many-cells results (**Fig. 4**), many exocrine cells that existed at Day1 disappeared at Day29 (the proportion of exocrine cells in total cells was decreased from about 44% to about 16% of total cell population). There were also non-pancreas specific cells such as stellate, endothelial, macrophage, mast, and Schwann cells that were present at Day1 but were mostly absent at Day29, hence, presumably most of them were eliminated during LTC. Interestingly, although there was a high percentage of endocrine cells that did not survive the LTC, all subtype cells of endocrine cells were present in sample of Day29. These data indicate that islets used for transplantation contain a substantial number of exocrine cells and other non-pancreatic cells, but endocrine cells can be cultured for a longer time than other types of cells (**Fig. 5-6**, **S6**). The loss of exocrine cells and non-pancreatic cells during culture may have unexplored impacts on the function of LTC-islets as showed in our cell-cell communication data below.

**Figure 6.**
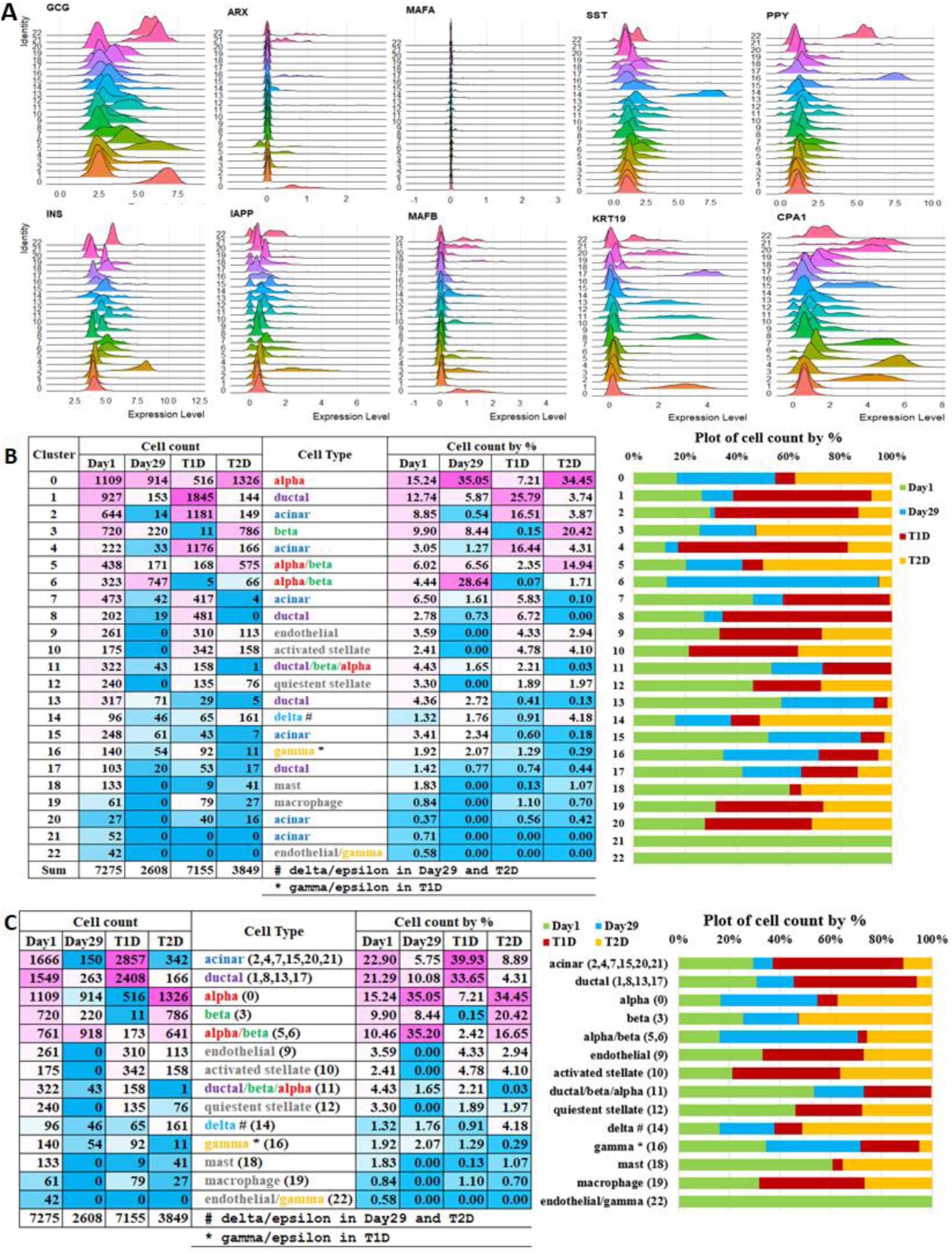
scRNA-seq analyses of LTC-islets—detailed analysis of cell clusters and cell types. **A.** Ridge plots of the expression ranges of marker genes of α-cells (*GCG, ARX, MAFB*), β-cells (*INS, IAPP, MAFA)*, δ cells (*SST*), γ cells (*PPY*), acinar cells (*CPA1*), and ductal cells (*KRT19*) in all predicted cell clusters. **B.** Detailed analysis of predicted cell clusters: the number of cells counts and percentages of cells counts in each cluster in each group of samples were listed on the left side and the percentage data were also plotted as bar graph on the right side. **C.** Detailed analysis of predicted cell types: the number of cells counts and percentages of cells counts in each cell type in each group of samples were listed on the left side and the percentage data were also plotted as bar graph on the right side.

### LTC-islets have a changed composition of cell subtypes

Despite all subtypes of cells in islets were still present at Day29, the composition of cell subtypes in islets at Day29 was changed compared to Day1, T_1_D, or T_2_D. Comparing cells from Day29 to Day1, α-cells had a much higher survival rate than β-cells. On Day1, the percentages of the largest groups of canonical α-cells (cluster-0) and canonical β-cells (cluster-3) were about 15% and 10%, respectively. On Day29, they were changed to about 35% and 8.5%, respectively. This reflects a change in the ratio of α-cell to β-cell from about 3 to 2 on Day1 to about 4 to 1 on Day29 (**Fig. 6B-C**, **S6, Table S2**). This resembles the α-cell to β-cell ratio under the T_1_D condition, but the selective loss of β-cells and the change of the α-cell to β-cell ratio in LTC is at a much lower intensity than in T_1_D (α-cell to β-cell ratio is more than 50 to 1 in T_1_D). In contrast, T_2_D has a similar ratio of α-cell to β-cell ratio as the sample of Day1. In addition to the loss of the majority of β-cells, T_1_D samples differ from all other samples by containing significantly fewer α-cells—a 50% reduction compared to the Day 1 sample—while having substantially more exocrine, endothelial, and stellate cells. Compared to the Day1 sample, both samples of Day29 and T_2_D have more α-cells and their numbers were more than doubled.

However, T_2_D samples still maintained their α-cell to β-cell ratio (**Fig. 6B-C**, **S6**). T_2_D samples have a high number of α-cells and β-cells but fewer acinar or ductal cells, which indicates some exocrine cells may be lost or transdifferentiated into endocrine cells in T_2_D. The major difference between sample of Day1 and other samples is the subgroup of cells, classified as either α-cells or β-cells, that formed cluster-5 and cluster-6. These cells are double positive for *INS* and *GCG* (*INS^+^GCG^+^*). While T_1_D samples had the lowest number of cluster-5 cells, T_2_D samples were enriched with these cells, and the Day29 sample and the Day1 sample had a similar percentage of them, indicating cluter-5 cells are vulnerable under T_1_D condition, increased under T_2_D condition, and most of them will survive LTC (**Fig. 6B-C**). Cluster-6 cells formed a distinct cluster primarily present in the Day 29 sample, clearly distinguishing it from the other samples (**Fig. 6B**). Therefore, we further characterized cells in the cluster-5 and cluster-6.

### Cells of Day29 were enriched with *INS* and *GCG* double positive cells and they were the major groups of cells that were different between Day29 and Day1

We plotted the expression of marker genes of α-cells (*GCG, ARX, MAFB*) and β-cells (*INS, IAPP, MAFA*) in each sample group, the expression of marker genes in each subtype of cells (left panels: violin plots in **Fig. 7**), the marker gene distribution in all cells (middle panels: UMAP plots in **Fig. 7**), and the marker gene expression ranges in each sample group (right panels: ridge plots in **Fig. 7**) or in each individual sample (**Fig. S7**), to compare their expression across sample groups. Expression of *INS* was almost absent in the T_1_D sample; a subgroup of cells with higher expression of *INS* was enriched in samples of T_2_D while cells with lower expression of *INS* were markedly reduced in T_2_D samples compared to the Day1 sample; *MAFA* expression was elevated in β-cells of the Day29 sample; *IAPP* expression was elevated in the Day29 sample and it had a lower expression level in both the samples of T_1_D and T_2_D; Cells with a high level of *GCG* expression were lost in all samples of T_1_D but this group of cells were enriched in the sample of Day29; *MAFB* expression was elevated in the α-cells of D29 and macrophage cells of T*_2_*D. *ARX* followed a similar expression trend as *GCG*. Hence, *IAPP* was highly expressed in β-cells and δ-cells of Day29, β-cells of Day29 were enriched with *MAFA* and are positive for both *MAFA* and *MAFB* (β-*MAFA^+^MAFB^+^-*cells), and α-cells of Day29 were enriched with *MAFB* and are positive for both *ARX* and *MAFB* (α-*ARX^+^MAFB^+^-*cells). These results agreed with the RNA-seq data that *IAPP*, *MAFA*, and *MAFB* genes were upregulated during LTC (**Fig. 1E-F**). These results led us to ask what could have caused β-cells and α-cells that survived LTC to have high expression levels of *MAFA* and *MAFB*, respectively? Another question is whether all β-cells and α-cells, or just subgroups of β-cells and α-cells that survived LTC have high *MAFA* and *MAFB* expression, respectively? To answer these questions, we looked at the subgroups of cells that were classified as either α-cells or β-cells, specifically, cells of cluster-0, -3, -5, and -6, in samples of Day1 versus Day29, and compared their presence in both samples of T_1_D and T_2_D (**Fig. 6B-C**, **8A-B**). Cells in cluster-0 are canonical mature α-cells because they express both *GCG* and *ARX* genes. These cells had a high level of *MAFB* expression in both Day1 and Day29 samples, but *MAFB* expression was reduced in both T_1_D and T_2_D samples (**Fig. 8A-B**). Hence, the loss of *MAFB* expression in mature α-cells could be a major indicator or a cause for both T_1_D and T_2_D, and restoring *MAFB* expression in mature α-cells may have therapeutic value for both T_1_D and T_2_D. Cells in cluster-3 are canonical mature β-cells because they express both *INS* and *MAFA* genes. *MAFA* expression was elevated in the sample of Day29 compared to the sample of Day1, and to some extent in the sample of T_2_D (**Fig. 8A-B**). Therefore, LTC led to higher *MAFA* expression in the Day29 sample for mature β-cells, indicating β-cells with a high *MAFA* expression level may be more fit for survival in LTC or the elevated *MAFA* expression may enable these cells to be more tolerant of harsh environments. Cells in cluster-5 are *INS^+^GCG^+^* cells and they were more like α-cells because they had a higher expression of *GCG*. *MAFB* expression was relatively higher in cells of cluster-5, and it also had higher expression in the sample of Day29 than that in the sample of Day1. On the other hand, *MAFA* had very low expression in these cells across all samples. Therefore, cells in cluster-5 were more like less mature α-cells, and LTC led to elevated *MAFB* expression in both mature and less mature α-cells, possible a critical reason for more α-cells, but less β-cells survived the LTC (**Fig. 6B-C**, **8A-B**). Cells in cluster-6 are the most interesting group of cells. They were more like β-cells as indicated by more cells in this cluster expressing *INS* and *IAPP*. Cluster-6 was also the largest group of cells that could distinguish the sample of Day29 from the samples in other three groups, but cells of cluster-6 had very low expression of both *MAFA* and *MAFB*. It is possible that cells in cluster-5 contain the original *INS^+^GCG^+^*cells that represent less mature α-cells and that later-appearing *INS^+^GCG^+^*cells in cluster-6 represent less mature β-cells which are supported by low *MAFA* or *MAFB* expression (**Fig. 6B-C**, **8A-B**). However, it is hard to tell whether the existing high percentage of cluster-6 cells in the sample of Day29 was caused by their higher survival rate than mature β-cells of cluster-3 during LTC, or if it was the result of transdifferentiation from other cells, such as cluster-5 cells, to β-cells during LTC. Lineage tracing data showed cells in cluster-5 were classified as cells that appeared in the earliest developmental stage, and cells in cluster-6 were classified as cells that appeared at a later time in early stages (**Fig. 8C-F**).

**Figure 7.**
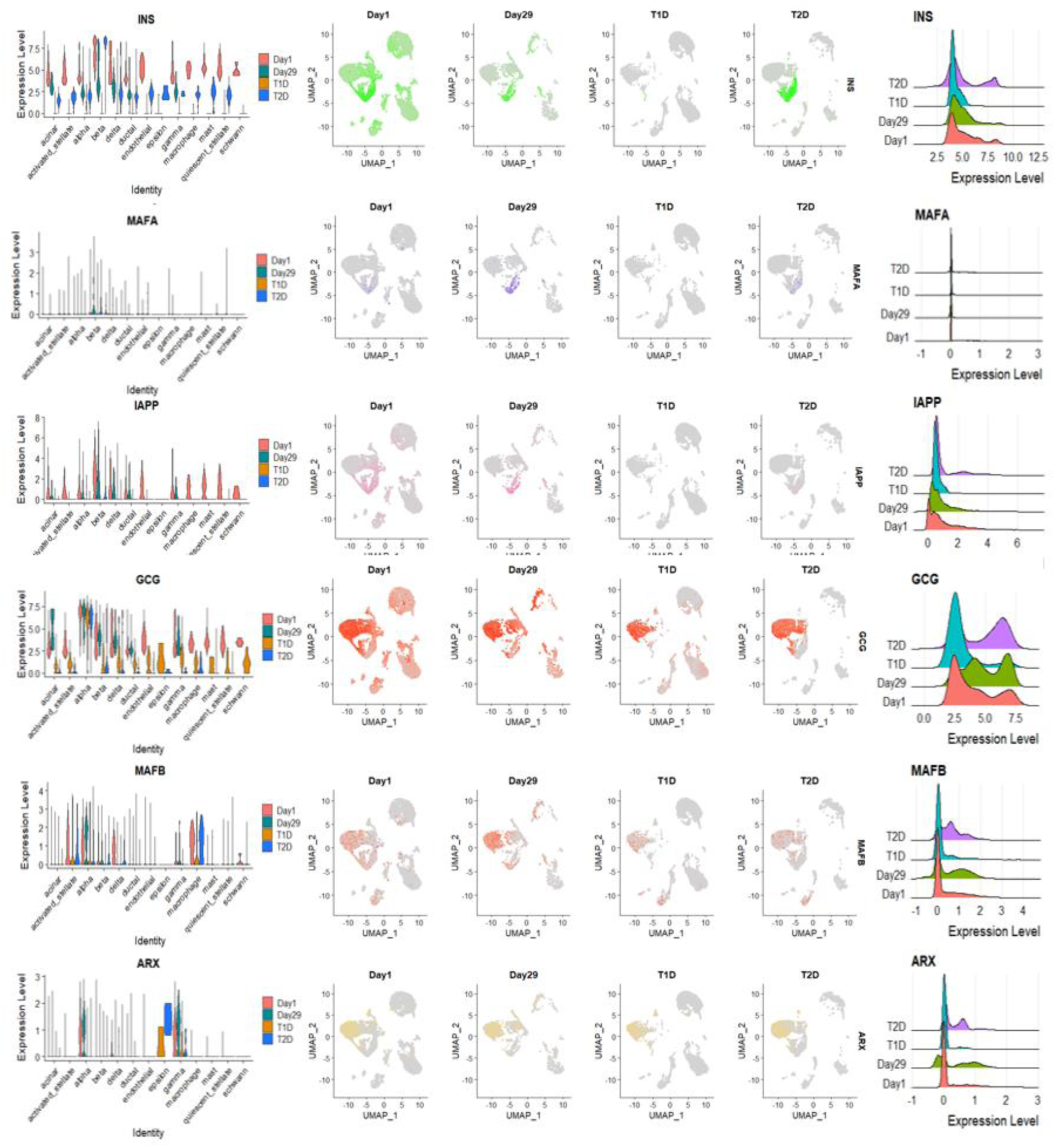
scRNA-seq analyses of LTC-islets—detailed analysis of the expression of marker genes of α-cells and β-cells in each group of samples. Left panels: side-to-side comparison violin plots to show the expression distributions of the six marker genes in each group of samples; Middle panels: UMAP plots to show the expression of the six marker genes in cells of each group of samples; Right panels: comparison ridge plots to show the expression range of the six marker genes in each group of samples.

**Figure 8.**
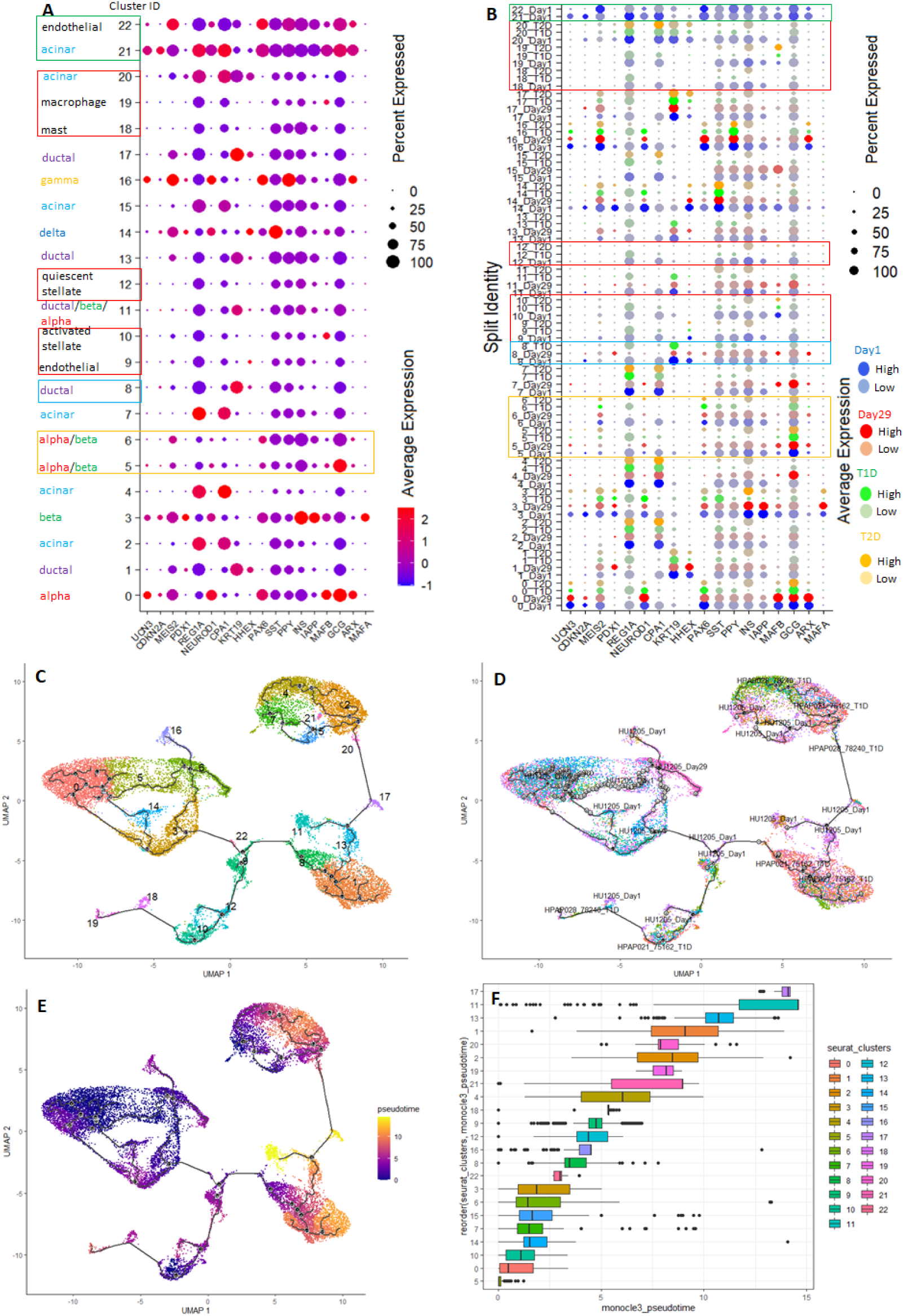
scRNA-seq analyses of LTC-islets—detailed analysis of gene expression in cell clusters and cell types. **A.** A dot plot showed the expressions of several selected genes (cell proliferation markers, major hormones, and transcription factors of pancreas) in clusters of cells and cell types. **B.** A dot plot showed the expressions of several selected genes in clusters of cells in each group of samples. **C.** The trajectory map of cells predicted by Monocle3, and nodes were labeled by predicted cell clusters. **D.** The trajectory map of cells predicted by Monocle3, and nodes were labeled by individual sample names. **E.** A heatmap plot showed Monocle3 predicted pseudotime of cells in all samples. **F.** A boxplot showed Monocle3 predicted pseudotimes of all cell clusters.

Therefore, the cluster-5 cells have some characteristics of pancreatic progenitor-like cells and there is a possibility that cluster-6 cells are derived from cluster-5 cells. Both cluster-5 and 6 cells express *MEIS2* and *PAX6* genes that are important for the development and function of endocrine cells ^28,29^ (**Fig. 8A-B**). Both samples of Day1 and Day29 have a similar percentage of cluster-5 cells, indicating that most cluster-5 cells may have survived the LTC (**Fig. 6B**). DEG analysis showed that cells in cluster-5 had a similar gene expression profile between Day29 and Day1 because there are only several significantly changed genes (**Supplementary file S1**). On the contrary, cluster-6 cells were mainly present in Day29, but had a low percentage in both the Day1 and T_2_D samples, and are almost absent in T_1_D samples, like β-cells of cluster-3, agreeing with the result that T_1_D patients usually loss majority of their functional β-cells, and only a small group of immature β-cells with α-cell characteristics were left in the residual β-cells of T_1_D ^30–32^ (**Fig. 6B-C**, **8A-B**). DEG analysis using too-many-cells data also revealed significant differences in pathways between N19 (cluster-3) and N13 (cluster-6) cells. Genes that are downregulated in N13 compared to N19 are involved in responses to nutrients such as lipids, organic cyclic compounds, calcium ions, and vitamins. This suggests that cluster-6 cells may be less sensitive to nutrient levels and have a higher likelihood of surviving LTC (**Fig. S8A-B**). Further in-depth analysis of DEGs between cell of cluster 6 and 3 showed that there are many common dysregulated genes between Day29 and Day1, in agreement with the too-many-cells results, and also showed that many pathways were altered after LTC by down-regulated or up-regulated genes. The differences between cells in cluster-6 and cluster-3 are much greater in the Day29 compared to Day1. This is evidenced by the fact that more pathways are altered in the Day29 sample, suggesting that LTC increases the divergence between cells in cluster-3 and cluster-6 (**Fig. S8C-F**).

### Sample of Day29 loses pancreatic progenitor cells or proliferating cells

By plotting the expression of cell proliferation markers, major hormones, and pancreatic transcription factors across cell clusters, we observed the following: cells in cluster-8 were specifically absent in T_2_D samples; clusters 21 and 22 were present only in the Day1 sample; and clusters 9, 10, 12, 18, 19, and 20, which consist of non-pancreatic cells, were absent in the Day29 sample. *UCN3*, the pancreatic progenitor marker, was mainly expressed in α-cells (cluster-0) and to some extent in β-cells and γ-cells (cluster-3 and cluster-16). The cell proliferation marker *Ki67* (*CDKN2A*) was mainly expressed in α-cells and β-cells. However, both *UCN3* and *Ki67* were expressed in acinar cells of cluster-21 which only presented in the Day1 sample, indicating that pancreatic progenitor or proliferating cells were either lost or differentiated into other cells during LTC; they were also lost in samples of T_1_D and T_2_D. Therefore, the loss of pancreatic progenitor or proliferating cells is a common problem for islets of LTC, T_1_D, and T_2_D (**Fig. 8A-B**).

### Ductal cells of Day29 expressed marker genes of endocrine cells

One group cells of interest are the cells in cluster-11, which express *KRT19*, *GCG*, and *INS*. They are classified as ductal cells, but also express major hormones secreted by both α- and β-cells, indicating that they might have the potential to transdifferentiate into α-cells or β-cells, and also supporting the idea that some ductal cells function as pancreatic progenitor cells ^33,34^.

Pseudo-time analysis also showed that cells in cluster-11 belong to cells at very late stages, suggesting that they may be in a transient stage and supporting the transdifferentiation from ductal cells to α or β-cells scenario^34^. However, samples of T_2_D lacked cluster-11, support an idea that reintroducing this group of cells to T_2_D patents may be a potential therapeutic strategy (**Fig. 8A-B**). Pseudo-time data also showed acinar and ductal cells are enriched in T_1_D samples with some of these cells appearing at very late stages, indicating that transdifferentiation of acinar and ductal cells to β-cells may occur in T_1_D ^34^ (**Fig. 8C-F**).

### LTC-islets have a changed cell-cell communications

To check whether the cell composition changes, and pathway changes will reflect on cell-cell interaction and function of islets, we performed CellChat analyses ^35^. Cell-cell communication analyses showed that both the number and strength of interactions experienced major changes among all four groups of samples. The interactions between endothelial cells and other types of cells, which mediated a large part of cell-cell communication in Day1 sample, was affected the most. This was lost in Day29 and was dramatically reduced in both T_1_D and T_2_D samples. In contrast, the communication between ductal and alpha cells in the Day29 sample, as well as between ductal and activated stellate cells in both T_1_D and T_2_D samples, stood out as the primary cell-cell interactions. In the Day 29 sample, ductal cells exhibited significant communication with alpha cells, and sent signals to both delta and gamma cells (**Fig. S9**). These data support a model where ductal cells play a critical role in the pancreas that they can communicate with other cell types and may transdifferentiate into other cell types.

## Discussion

In both diabetes research and therapeutic developments, *in vitro* culture of isolated human islets is required, and human islets are precious due to the limited supply of donor pancreases. *In vitro* culture offers several potential benefits for cultured islets, including the removal of immune cells that could cause immunotoxicity, the opportunity for application of reagents to enhance islet functionality, and additional time to pool islets from different donors, assess islet quality before grafting, and prepare them for transplantation ^4^.

While islets for transplantation only need to be cultured for a short time, islets for research may need to be cultured for a much longer time. However, islets undergoing *in vitro* culture usually result in a loss of cell viability and function. There are some intriguing questions, such as, what is the *in vitro* culture time-limit for islets to be functional, which cells are lost and whether the cell loss is a stochastic or a selective process, and which genes are shut down or activated during islet *in vitro* culture. Answering these questions may aid in the development of novel procedures to maintain islets in culture for an extended time frame without significant loss of cell viability and function, to engineer islets with enhanced or novel functions, and to better interpret experimental results using *in vitro* cultured human islets.

Both our RNA-seq and smRNA-seq results showed LTC-islets went through major gene or miRNA expression changes and many cellular pathways were affected. The altered genes or miRNAs are mainly involved in the extracellular matrix, insulin secretion, cell proliferation, cell adhesion, innate immune responses related to focal adhesion, the PI3K-AKT signaling pathway, ECM-receptor interaction, and the RAP1 and MAPK signaling pathways (**Fig.2**). Our smRNA-seq results also implied cellular antifungal functions may be triggered in LTC-islets which maybe a potential problem for *in vitro* islet culture. Our scRNA-seq data showed, after LTC, cell compositions and cell-cell interactions were changed. Epithelial cells and other nontypical pancreatic cells were eliminated, and both acinar cells and ductal cells of exocrine cells were largely reduced. The reduction of exocrine cells may have major impacts on the function of LTC-islets since cell-cell interactions were altered and previous studies showed lower purity of islets that contain exocrines cells have a better transplantation outcome ^36^. Most islet cell subtypes survived LTC with α-cells exhibiting a much higher survival rate than β-cells. As a result, the β-cell to α-cell ratio was significantly reduced, resembling the phenomenon observed in T_1_D. β-cells that survived LTC also exhibited a very different gene signature from β-cells in islets cultured for a short time. This resembles the residual β-cells in T_1_D that have a different gene expression profile from normal β-cells. However, β-cells of LTC and residual β-cells of T_1_D are different in several important aspects. For instance, the residual β-cells of T_1_D display premature features and lack normal β-cells function, which may have mediated their escape from the autoimmune attack ^37^. In contrast, β-cells in LTC contained both mature and immature cells with many expressing both *MAFA* and *MAFB* genes, making some of them β-*MAFA^+^MAFB^+^-* cells.

Our data supports the idea that cell death in islets under culture conditions is not random. Instead, it reflects a selective survival process, where certain cell subtypes within the islet exhibit greater fitness and are less vulnerable to elimination than others. Our results showed that α-cells have a higher survival rate than β-cells, and most β-cells are also more vulnerable to cell death in LTC. The higher death rate of β-cells than α-cells under the same LTC condition suggests that there are intrinsic mechanisms in β-cells that contribute to their death, while mechanisms in α-cells may confer resistance to cell death in unfavorable environments. It has been reported that human islets are heterogeneous and contain four distinct subtypes of β-cells ^25^. Follow-up studies showed α-*ARX^+^MAFB^+^-*cells and β-*MAFA^+^MAFB^+^-*cells are subpopulations of α-cells and β-cells exist in native pancreatic tissue. It also showed that the subgroups of α- or β-cells expressing two transcription factors exhibit a greater expression of key genes related to glucose sensing and hormone secretion relative to other subpopulations that express only one or neither transcription factor ^26^. The two transcription factors in α-cells or β-cells may act cooperatively to make them highly functional mature subpopulations. Hence, both α-cells or β-cells that survived LTC may be cells derived from these highly functional, mature subpopulations, and those less mature subpopulations may be easily eliminated during LTC, probably due to their poor ability for glucose metabolism. In our study, however, the cluster-6 cells were β-*MAFA^+^MAFB^+^-*cells and they were mainly present in the Day29 sample. It is an interesting question whether these cells are immature β-cells with a higher survival rate in LTC, transdifferentiated cells from α-cells, or dedifferentiated β-cells that sacrificed function for survival ^37^. They were present at a similar percentage as mature α-cells (28% versus 35%) and make up the 2^nd^ largest cell population in the Day29 sample. This contrasts with the result that most residual β-cells from T_1_D were immature β-cells that lacked normal β-cells functions, as they were also present at a lower percentage. It is interesting that this group of cells was also elevated in T_2_D samples ^30–32^. These data support the idea that, in addition to immune attacks, there are other factors contributing to β-cells death in T_1_D ^38^. Hence, this raises the question of which factor contributes the most to β-cell death in different individuals of T_1_D: autoimmune attacks on the pancreas, or the stress conditions faced by the pancreas during the development of T_1_D.

One limitation in our study is that we have used the 2-D culture system for LTC which is simple and easy to handle, not the more sophisticated 3-D system to mimic the *in vivo* living environment. It is not surprising that the extracellular matrix of islets cells, disrupted under *in vitro* culture, has a major change in affected pathways. However, even in a 3-D *in vitro* culture system, the living condition will still be far from an *in vivo* environment. LTC-islets will face many unfavorable factors, creating a stress-inducing environment.

The other limitation in our study is that we do not have single cell smRNA-seq data to associate DEmiRs with the cell composition of residual cells. Our RNA-seq data and scRNA-seq data showed cell population change in islets is associated with DEGs. Compared to protein-coding genes, miRNAs make up less than 10% of the total. The number of essential or conserved miRNAs is only a few hundred, and the number of highly expressed miRNAs in a particular tissue or cell is even smaller. Importantly, miRNAs generally act as post-transcriptional regulators of gene expression. All these features make miRNAs ideal differentially expressed genetic markers. We observed that major changes in miRNAs happened earlier than in genes. Most miRNAs were downregulated by day seven in LTC while most genes were downregulated within two weeks in culture. The significantly downregulated miRNAs outnumbered the significantly upregulated miRNAs by more than a 2 to 1 ratio (81 versus 29). The three most abundant miRNAs, miR-375-3p, miR-7-5p, and miR-148a-3p were all altered. While miR-375-3p was upregulated, both miR-7-5p and miR-148a-3p were downregulated during LTC. The changes of the above three miRNAs correlated well with results that there are more miR-375-3p and miR-7-5p-enriched endocrine cells than miR-148a-3p-enriched exocrine cells that survived the LTC. The increase of miR-375-3p abundance corresponded with a change in cell ratio of α-cells to β cells. Because most of the dysregulated miRNAs are not miRNAs enriched in α-cells or β-cells according to published miRNA profiling data ^16^, we are not able to predict whether those upregulated miRNAs are enriched in α-cells and those downregulated miRNA are enriched in β-cells. This is especially challenging as many cell subtypes disappeared during LTC, making it even more difficult to determine the cause of downregulated miRNAs. Nevertheless, the upregulated miRNAs should be miRNAs enriched in α or β-cells since most islet cells surviving LTC are α or β-cells, or they may be miRNAs that can mediate islets cell survival in LTC. It is possible some of these dysregulated miRNAs can be used as markers to access the quality of LTC-islets or to track the performance of transplanted islets.

In summary, our gene and miRNA profiling data and scRNA-seq analysis data showed gene and miRNA expression level and cell composition changed in LTC-islets. These data may represent a valuable resource to support the creation of novel islets culture protocols to expand islet culture time while preserving islet function and may guide us to genetically engineer human islets to maintain their viability. Some differentially expressed genes, especially miRNAs in LTC-islets, may be used as markers to assess the quality of islets and as markers to indicate the success or failure of islets transplantations. Our observation of cluster-5 and cluster-6 cells indicated LTC-islets could be used to study human islet cell transdifferentiation or dedifferentiation *in vitro*, a process that has been largely limited to research using samples from T_1_D and T_2_D patients. One approach to bypass the limitation of using human pancreas from cardiac donors is to use stem cell derived islet-like cells for diabetes therapy or research.

However, although these cells can perform major functions of β-cells, they are not fully functional as primary islets. Possible causes include incomplete maturation of endocrine cells, incorrect cell ratios, and improper spatial arrangement of cells. Our cell-cell interactions results suggest that other cell types, such as endothelial and ductal cells, are critical for the survival and function of islets. In that sense, the results in our study may help to create novel protocols for generation of stem cell-based islet-like cells with improved function. Our scRNA-seq data was limited to a single sample. Future scRNA-seq should be performed on multiple samples and at various culture time points to further explore LTC-islets. This approach will help identify the optimal LTC time point for isolating specific cell populations, such as those in cluter-5 and cluster-6, for functional studies. In addition, we think pancreases that are not qualified for transplantation may be submitted to LTC to eliminate low viability cells, and to collect high viability cells that can re-aggregate into islets with higher quality.

## Materials and Methods

### Isolation of human pancreatic islets

Pancreata from brain-dead cadaveric donors were provided by OneLegacy (Los Angeles, CA), stored in preservation solution after flushing with cold University of Wisconsin preservation solution and were transported on wet ice to a Good Manufacturing Practices facility at City of Hope National Medical Center (Duarte, CA) for processing. Pancreatic islets isolation protocol was followed as described^15,39^. Briefly, pancreata were digested using collagenase supplemented with either thermolysin or neutral protease. Then, the digested pancreatic tissue was collected, washed, and purified in a cooled COBE 2991 cell processor (COBE Laboratories Inc., Lakewood, CA, USA. The islets were cultured at 37°C with 5% CO_2_ for 24-72 hours before assessing purity. The use of human pancreatic tissues was approved by the Institutional Review Board of City of Hope National Medical Center (IRB#01046). Only organs from nondiabetic individuals (HbA1c <6.5%) were included. Informed consent was obtained from the legal next of kin of each donor. Deidentified donor demographic information is summarized in Supplementary Table S1.

### Human islets culture

Human islets were cultured in the same medium as islets were used for transplantation^15^. Briefly, islets were cultured in Connaught’s Medical Research Lab (CMRL)1066 medium (Corning, 98-304-CV) supplemented with 0.5% human serum albumin (Baxalta, NDC 0944-0493-01) and IGF-1 (0.1 µg/mL) (Repligen, 10-1011-125) for less than one day after isolation from the pancreas before being dissociated into single cells for LTC or scRNA-seq. For RNA-seq and smRNA-seq, an aliquot of isolated islets cultured for less than 24 hours were used to isolate total RNAs as day1 (D1) samples, then the rest of the islets were digested with Gibco’s gentle cell dissociation reagent TrypLE (ThermoFisher, Waltham, MA) and cultured for one to four weeks to obtain samples at different time points in culture. Every three days, cells were briefly spun down and resuspended in fresh medium. An aliquot of cultured islets was taken out every week to isolate total RNAs as samples for day7 (D7), day14 (D14), day21 (D21), and day28 (D28). For scRNA-seq, about 10,000 cells were used for scRNA-seq on the same day right after dissociation and 7275 cells produced good quality sequencing results after filtering (Day1). About 50,000 dissociated cells were cultured for four weeks. Every three days, cells were briefly spun down and resuspended in fresh medium. On the final day, residual cells were dissociated for scRNA-seq (Day29) and 2348 cells gave good quality sequencing results after filtering.

### RNA isolation

Total RNA was isolated from islets using TRIzol (ThermoFisher, Waltham, MA). The manufacturer’s instructions were followed throughout. The RNA integrity and quality was determined, and amounts were quantified using a Qubit 4, a Nanodrop, and an Agilent Bioanalyzer.

### RNA deep sequencing, data processing, analysis, and statistics

RNA deep sequencing was carried out as previously published ^40^. Briefly, 100 ng to 1.0 µg of RNA was used to construct RNA-seq libraries for single reads, flow cell cluster generation, and 51 cycle (51-nt) sequencing on a HiSeq 2500 (Illumina, San Diego, CA). The sequence depth was 30M reads per sample with 8 samples per lane (by barcoding), and was performed by the Integrative Genomics Core of City of Hope National Medical Center. Raw reads obtained from RNA-seq were mapped to the human hg38 genome (iGenomes from Illumina) using the subread aligner (version 2.0.6) and read counts of genes was obtained with featureCounts. Subread aligner and featureCounts in Subread package (release 2.0.6) was used with default parameters ^41^. Raw read count from featureCounts were input into Bioconductor package DEseq2 (version 1.40.1) in R version 4.3.0 to generate the differential expressed gene results at the default setting [the Wald-test was applied to assess the *p* value for differential gene expressions and the adjusted *p* value (*p-adj*) was done by the Benjamini and Hochberg method]^42^. Unless specified in the figure legends, p-adj < 0.05 was used as cutoff for gene expression with significant changes.

### Small RNA deep sequencing, reads processing, and data analysis

Small RNA deep sequencing was carried out using a customized protocol ^15,43,44^. Briefly, 100 ng to 1.0 µg of RNA was used to construct small RNA libraries. The 5′ adaptor used in the Illumina (San Diego, CA, USA) Truseq small RNA deep sequencing kit was replaced with a customized 5′ adaptor by adding 3 random nucleotides (nts) at the 3′ end of the original 5′ adaptor sequence to increase complexities of PCR amplifications and reduce biases caused by both PCR and adaptor ligation. The 3′ adaptor provided in the kit was used for the 3′ end ligation and sample barcoding. Single reads, flow cell cluster generation, and 51 cycle (51-nt) sequencing was performed on a HiSeq 2500 (Illumina, San Diego, CA, USA) by the Integrative Genomics Core of City of Hope National Medical Center. Raw smRNA-seq reads for each sample were separated by barcodes. Then, the 5’ adaptor with the three random nts was removed by Cutadapt prior to further processing and analyzing using miRge2 ^45,46^. Unnormalized miRNA read counts generated by miRge were used to calculate DEmiRs by DEseq2. The same procedure for calculating DEGs in RNA-seq was followed.

### Single cell RNA sequencing and data analysis

scRNA-seq was performed jointly by the Integrative Genomics Core (construction of libraries and quality check of sequencing runs) and the Translational Genomics Research Institute (sequencing) of City of Hope National Medical Center. After islets dissociation using TrypLE (ThermoFisher, Waltham, MA), dead cells were removed by Dead Cells Removal Microbeads (Miltenyl Biotec, Bergisch Gladbach, Germany), and cell viability was checked. Samples with cells at >80% viability were loaded onto the Chromium Controller (10X Genomics, Pleasanton, CA, United States) targeting 5,000–10,000 cells per reaction for the construction of libraries. The Chromium v3 single-cell 3′ RNA-seq reagent kit (10X Genomics) was used to generate scRNA-seq libraries according to the manufacturer’s protocol. The libraries were sequenced using the NovaSeq 6000 system (Illumina, San Diego, CA, United States) with a depth of 50k-100k reads per cell. Raw sequencing data were demultiplexed to fastq files using the Cell Ranger (version 6, 10X Genomics) and aligned to the human hg38 genome. The filtered matrix data generated by Cell Ranger were used for the next step data analysis using too-many-cells ^21^ and Seurat (version 4) ^24^.

We followed the example procedure of too-many-cells on github to analyze our scRNA-seq data. Then, performed integrated scRNA-seq data analysis on our scRNA-seq datasets with downloaded patients’ scRNA-seq datasets of T_1_D (GSE148073) ^22^ and T_2_D (GSE816085) ^23^ using the integrated scRNA-seq data analysis workflow provided in Seurat v4 ^24^. Briefly, the count data from low quality cells with <200 genes, < 500 transcripts were excluded. We predicted cells in all four datasets in our analysis (Day1, Day29, T_1_D, and T_2_D) according to the annotated cell types in published scRNA-seq datasets of the human pancreas (panc8) that were curated in SeuratData. Specifically, datasets of celseq, celseq2, smartseq2, and indrop, the datasets with more sequenced cells among all eight datasets were used as a reference for prediction cell types in our datasets ^24^. Next, the four datasets with predicted cell types were integrated and standard workflow post integration in Seurat was followed: the integrated data was scaled (ScaleData), a principal component (PC) analysis was performed using the top 30 PCs (RunPCA: npcs=30), UMAP dimension reduction was performed, the k-nearest neighbors of each cell were determined, and shared nearest-neighbor (SNN) graphs were constructed (FindNeighbors), then the clusters were identified using the SNN modularity optimization-based clustering algorithm (FindClusters, resolution = 0.5). Uniform manifold approximation and projection dimensionality reduction on the first 20 principal components, annotation of identified clusters, and generation of plots were performed by Seurat to visualize the clusters, heatmaps, and feature plots.

### Cell trajectory, developmental pseudotime, and cell-cell communication

Single-cell trajectory and pseudotime analysis were conducted by Monocle3 using the variable features identified by Seurat. A heatmap of the cluster-specific genes along the pseudotime was also generated using Monocle3 ^47^. Cell-cell communication was performed using CellChat R package. Standard analysis workflow in CellChat tutorial were followed ^35^.

### Other computational resources

R (version 4.3/4.4), R studio, and Bioconductor software and associated packages were used for data analysis and visualization. ComplexHeatmap ^48^, EnhancedVolcano ^49^, ggplot2, and pheatmap were used to generate heatmap plots and volcano plots. ClusterProfiler^50^, Enrichment plot, and ClusterGVis R packages were used for GO and KEGG analysis and results visualization plots.

## Statistical Analyses

Inferential statistics of differential gene or miRNA expressions in islets cells were calculated by the default setting in DESeq2. Descriptive statistics in gene/miRNA expression data or comparison of one gene/miRNA in two different data sets were calculated by two tailed T-tests. Unless specified in the figure legends, p-adj < 0.05 for DESeq2 data or p-value < 0.05 for others was used as the cutoff for significance in gene expression. Error is expressed as SD.

## Data Availability

All raw and some processed sequencing data generated in this study was submitted to the NCBI Gene Expression Omnibus (GEO; https://www.ncbi.nlm.nih.gov/geo/) under accession number GSE280983 (RNA-seq), GSE280940 (miRNA-seq), and GSE280984 (scRNA-seq).

## Acknowledgements

The City of Hope Core Facilities are supported by the National Cancer Institute of the National Institutes of Health under award number P30CA33572.

## Author contributions

G.S., M.Q., Y.S. and A.D.R. designed the experiments. OS, EL, and DH contributed to experiments, data analysis, and manuscript; G.S. performed most bioinformatic data analysis. G.S. and Y.S. wrote the manuscript with input from the other authors. All authors read and approved the final manuscript.

## Materials and correspondence

Correspondence and requests for materials should be addressed to G.S.

## Supplementary information

Supplementary information is available for this paper at https://

