## Supplementary data and file for "Multi-omics analysis of long-term cultured human islets"

### 1.Supplementary Tables

**Table S1.** Donors' demographic information and pancreases and islets processing details

| Donor ID | Age, year | Sex | Race | BMI, kg/m2 | HbA1c % | Cause of death | Islet purity % | Islet viability % | OPO | Total culture time, day |
|---|---|---|---|---|---|---|---|---|---|---|
| Hu1181* | 28 | M | Caucasian | 30.75 | 5.20 | HT | 85 | 98.7 | OneLegacy | 7, 14, 21,28 |
| Hu1192* | 31 | M | Hispanic | 37.37 | 5.00 | HT | 85 | 97.0 | OneLegacy | 7, 14, 21,28 |
| Hu1193* | 30 | F | Caucasian | 30.78 | 5.10 | HT | 80 | 98.0 | OneLegacy | 7, 14, 21 |
| Hu1205** | 42 | F | Caucasian | 31.15 | 5.50 | CVA | 80 | 94.0 | OneLegacy | 1, 29 |

*Bulk RNA-seq and smRNA-seq; ** scRNA-seq; BMI: body mass index; HbA1c: hemoglobin A1c; HT: Head trauma; CVA: Cerebrovascular accident; OPO: Organ Procurement Organization.

**Table S2.** Summary tables of detailed analysis results of predicted cell clusters versus predicted cell types.

A. Pancreatic specific cells

| Cluster | Alpha | Beta | Delta | Gamma | Epsilon | Acinar | Ductal |
|---|---|---|---|---|---|---|---|
| 0 | 3825 | 34 | 0 | 4 | 0 | 0 | 0 |
| 1 | 0 | 0 | 0 | 0 | 0 | 2 | 3065 |
| 2 | 0 | 1 | 1 | 0 | 0 | 1985 | 0 |
| 3 | 122 | 1564 | 42 | 4 | 0 | 3 | 0 |
| 4 | 11 | 1 | 1 | 0 | 0 | 1584 | 0 |
| 5 | 1089 | 247 | 11 | 1 | 0 | 3 | 0 |
| 6 | 684 | 424 | 27 | 2 | 0 | 0 | 0 |
| 7 | 39 | 18 | 0 | 1 | 0 | 878 | 0 |
| 8 | 10 | 1 | 0 | 0 | 0 | 0 | 691 |
| 9 | 16 | 55 | 2 | 0 | 0 | 0 | 15 |
| 10 | 0 | 0 | 0 | 0 | 0 | 0 | 0 |
| 11 | 56 | 107 | 12 | 1 | 2 | 15 | 331 |
| 12 | 13 | 59 | 1 | 0 | 0 | 4 | 1 |
| 13 | 8 | 28 | 8 | 0 | 0 | 6 | 372 |
| 14 | 18 | 29 | 317 | 0 | 1 | 0 | 3 |
| 15 | 20 | 40 | 4 | 0 | 0 | 255 | 0 |
| 16 | 10 | 17 | 3 | 259 | 7 | 1 | 0 |
| 17 | 2 | 2 | 1 | 0 | 0 | 6 | 182 |

| | | | | | | | |
|---|---|---|---|---|---|---|---|
| 18 | 0 | 12 | 1 | 0 | 0 | 0 | 1 |
| 19 | 1 | 5 | 0 | 0 | 0 | 2 | 1 |
| 20 | 0 | 0 | 0 | 0 | 0 | 77 | 6 |
| 21 | 10 | 0 | 0 | 0 | 0 | 42 | 0 |
| 22 | 5 | 0 | 0 | 19 | 0 | 0 | 2 |

B. Non-pancreatic specific cells

| Cluster | Endothelial | Macrophage | Mast | Activated stellate | Quiescent stellate | Schwann |
|---|---|---|---|---|---|---|
| 0 | 0 | 0 | 2 | 0 | 0 | 0 |
| 1 | 0 | 0 | 2 | 0 | 0 | 0 |
| 2 | 0 | 0 | 1 | 0 | 0 | 0 |
| 3 | 0 | 0 | 0 | 2 | 0 | 0 |
| 4 | 0 | 0 | 0 | 0 | 0 | 0 |
| 5 | 0 | 0 | 0 | 1 | 0 | 0 |
| 6 | 0 | 0 | 0 | 0 | 3 | 1 |
| 7 | 0 | 0 | 0 | 0 | 0 | 0 |
| 8 | 0 | 0 | 0 | 0 | 0 | 0 |
| 9 | 582 | 0 | 0 | 0 | 9 | 5 |
| 10 | 2 | 0 | 0 | 667 | 6 | 0 |
| 11 | 0 | 0 | 0 | 0 | 0 | 0 |
| 12 | 0 | 0 | 0 | 75 | 247 | 51 |
| 13 | 0 | 0 | 0 | 0 | 0 | 0 |
| 14 | 0 | 0 | 0 | 0 | 0 | 0 |
| 15 | 0 | 0 | 0 | 0 | 0 | 0 |
| 16 | 0 | 0 | 0 | 0 | 0 | 0 |
| 17 | 0 | 0 | 0 | 0 | 0 | 0 |
| 18 | 0 | 5 | 164 | 0 | 0 | 0 |
| 19 | 0 | 128 | 30 | 0 | 0 | 0 |
| 20 | 0 | 0 | 0 | 0 | 0 | 0 |
| 21 | 0 | 0 | 0 | 0 | 0 | 0 |
| 22 | 15 | 0 | 1 | 0 | 0 | 0 |

**Figure S1**: Volcano plots of DEGs between Day 7 (D7), Day 14 (D14), Day 21 (D21), Day 28 (D28), and Day 1 (D1). DEGs were plotted as log2 fold change ($log_2FC$) versus negative log10 pvalues ($-log_{10}P$).

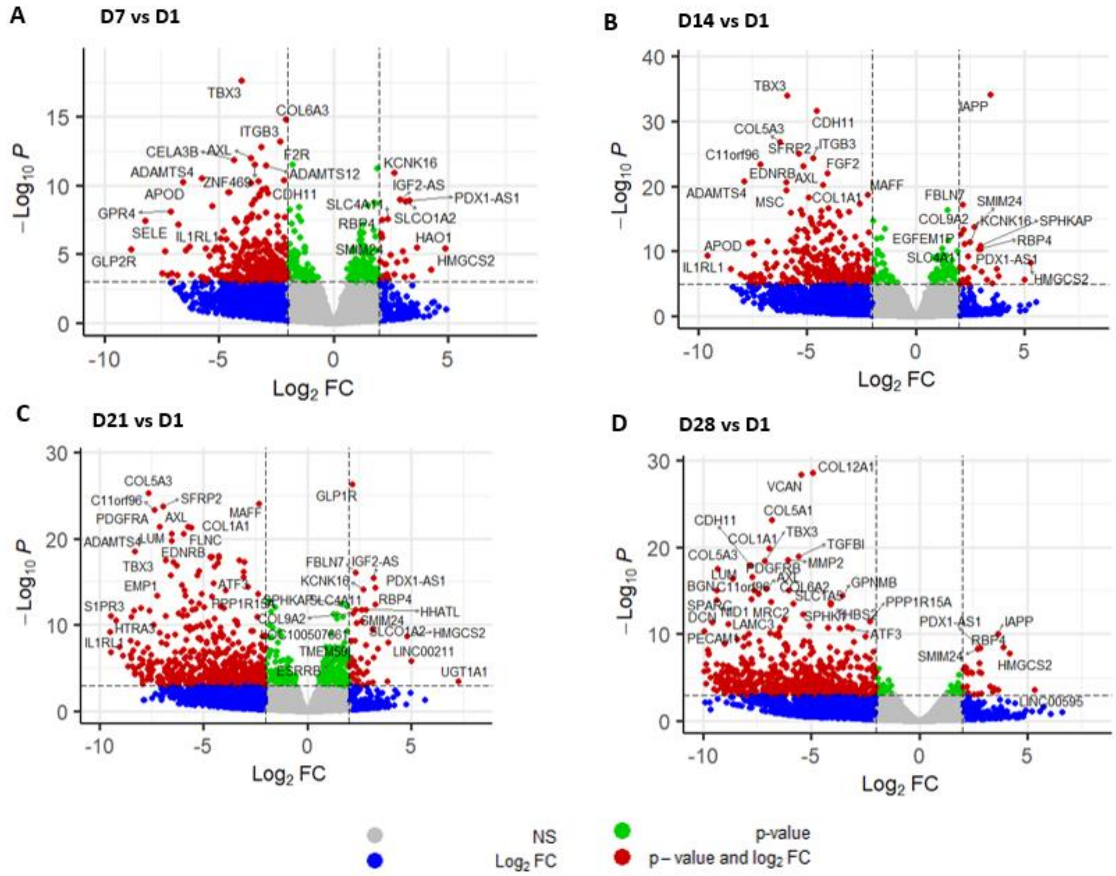

**Figure S2: GO and pathway analyses of the two upregulated DEmiRs in LTC-islets:miR-877-5p and miR-423-5p. A.** A bar plot of top-10 enriched GO terms and pathways. **B.** A bubble plot of top-10 enriched GO terms and pathways.

**A.**

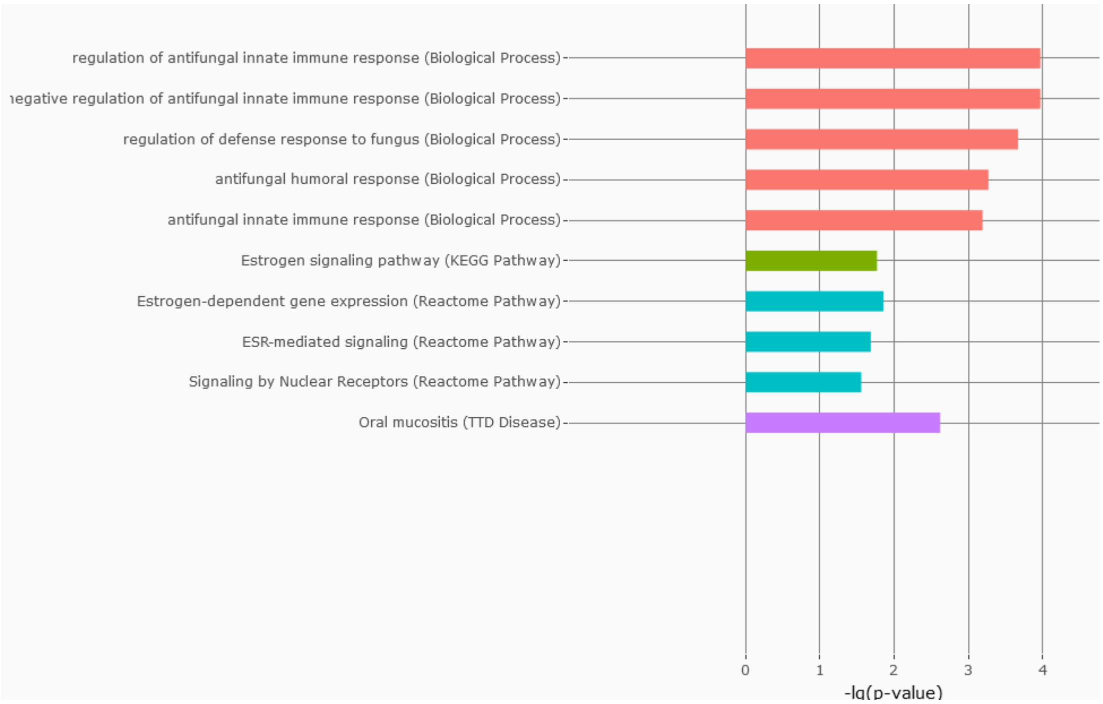


**B.**

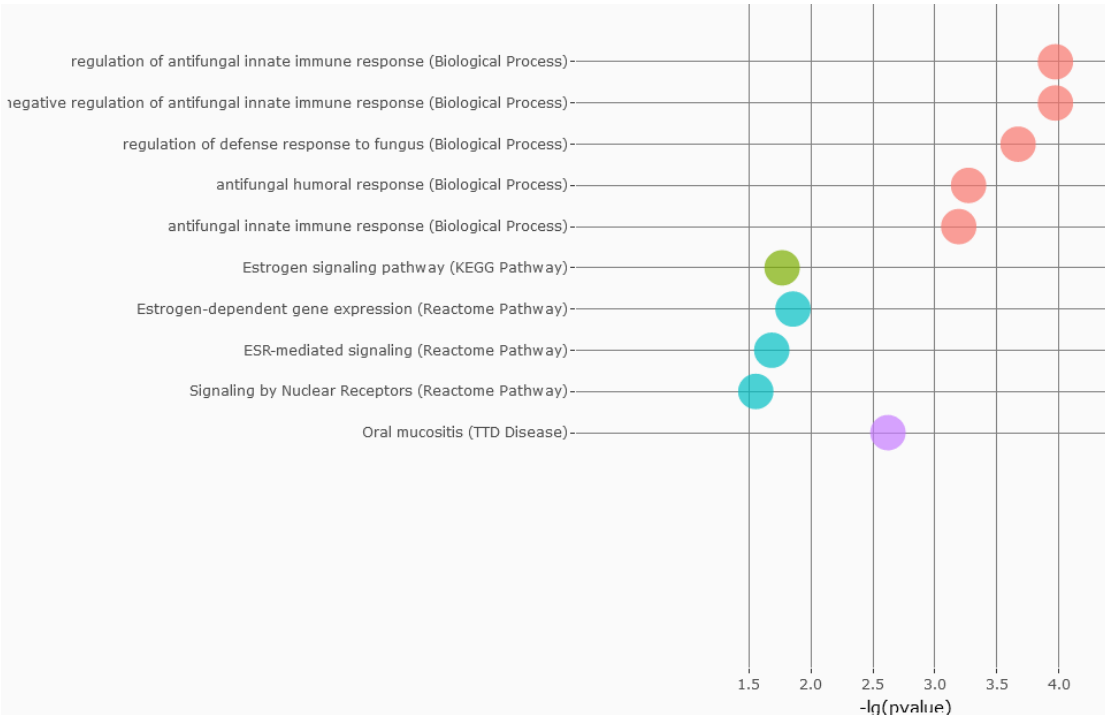

**Figure S3**: **GO and pathway analyses of the downregulated DEmiRs in LTC-islets. A.** A bar plot of top-5 enriched GO:BP, GO:CC, and KEGG pathways. **B.** A bubble plot of top-5 enriched GO:BP, GO:CC, and KEGG pathways. **C.** Protein-protein interactions formed by the top-50 genes targeted by the downregulated DEmiRs in LTC-islets. Interactions between IRS1 and IGF1R with others stood out.

**A.**

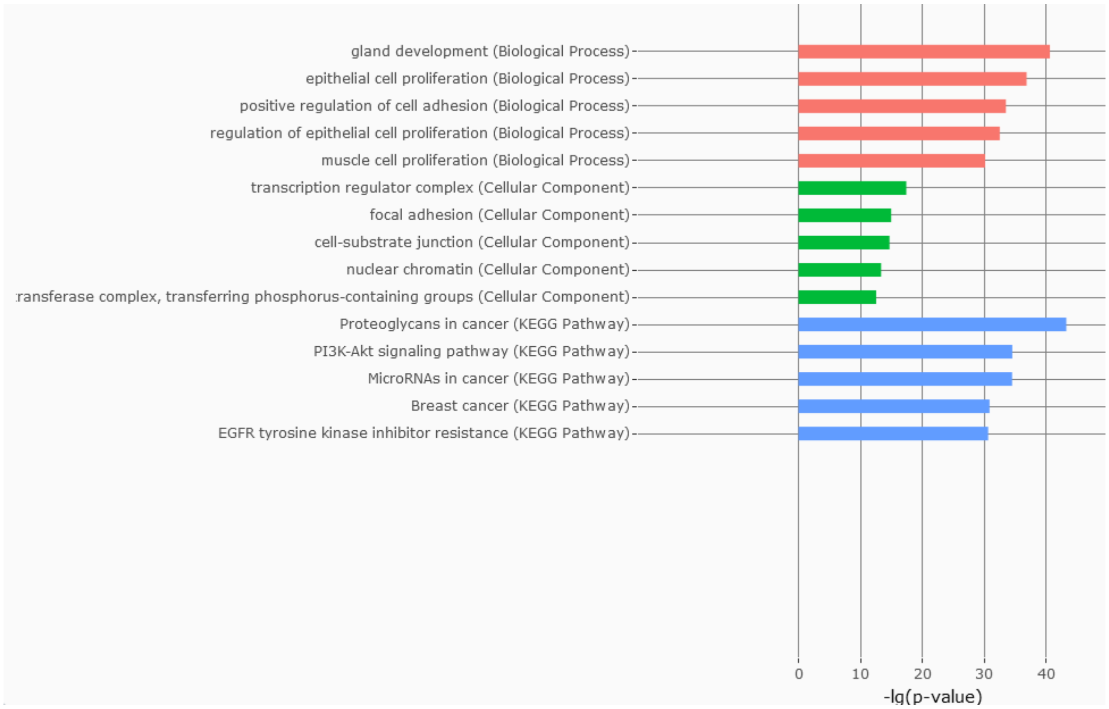


**B.**

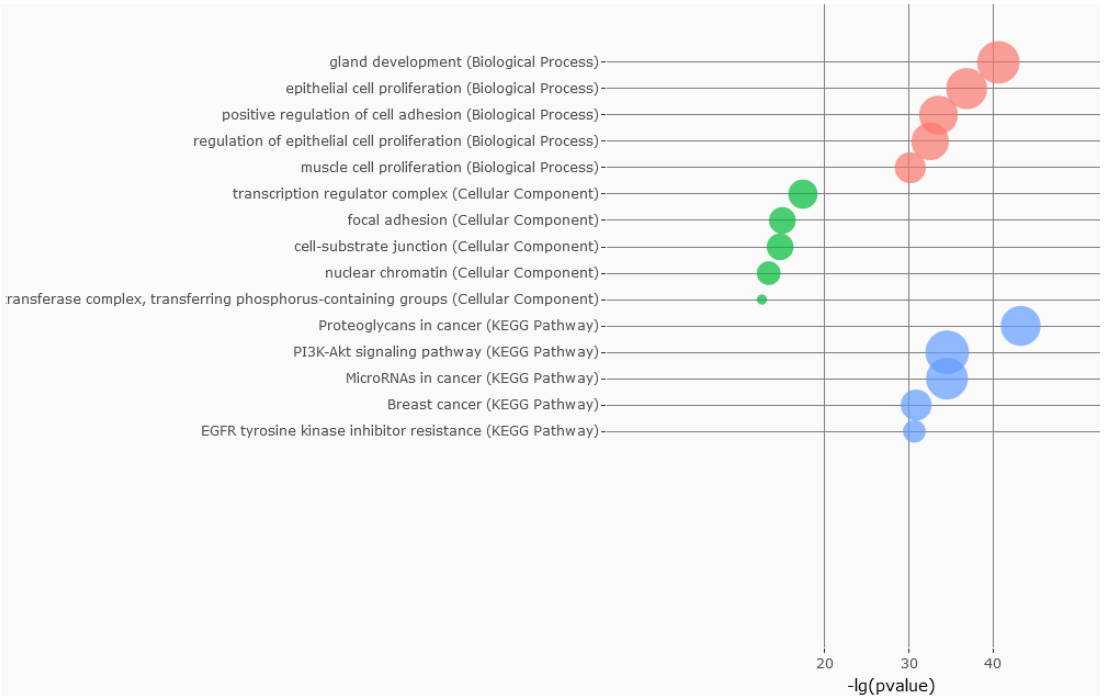
gland development (Biological Process)
epithelial cell proliferation (Biological Process)
positive regulation of cell adhesion (Biological Process)
regulation of epithelial cell proliferation (Biological Process)
muscle cell proliferation (Biological Process)
transcription regulator complex (Cellular Component)
focal adhesion (Cellular Component)
cell-substrate junction (Cellular Component)
nuclear chromatin (Cellular Component)
ransferase complex, transferring phosphorus-containing groups (Cellular Component)
Proteoglycans in cancer (KEGG Pathway)
PI3K-Akt signaling pathway (KEGG Pathway)
MicroRNAs in cancer (KEGG Pathway)
Breast cancer (KEGG Pathway)
EGFR tyrosine kinase inhibitor resistance (KEGG Pathway)
20
30
40
-lg(pvalue)

**C.**

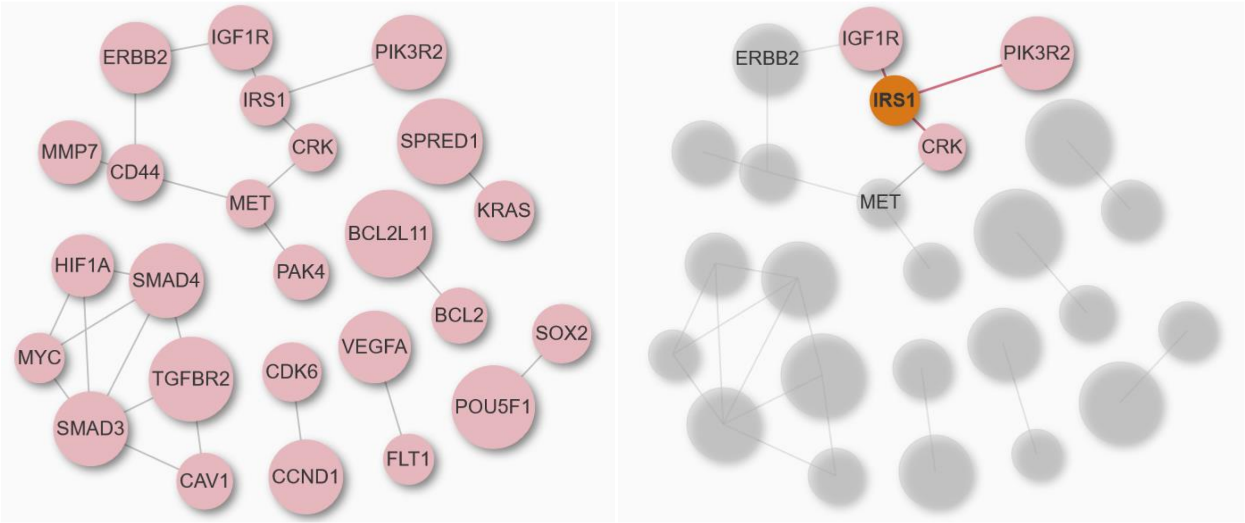
ERBB2
IGF1R
PIK3R2
IRS1
MMP7
CD44
CRK
SPRED1
MET
KRAS
BCL2L11
HIF1A
SMAD4
PAK4
BCL2
SOX2
MYC
TGFBR2
CDK6
VEGFA
POU5F1
SMAD3
CAV1
CCND1
FLT1
ERBB2
IGF1R
PIK3R2
IRS1
CRK
MET

**Figure S4**: **A volcano plot of DEGs between $T_1D$ and healthy individuals.** RNA-seq dataset of $T_1D$ (GSE162689 from GEO database) was downloaded and reanalyzed for DEGs of $T_1D$ versus healthy individuals (normal).

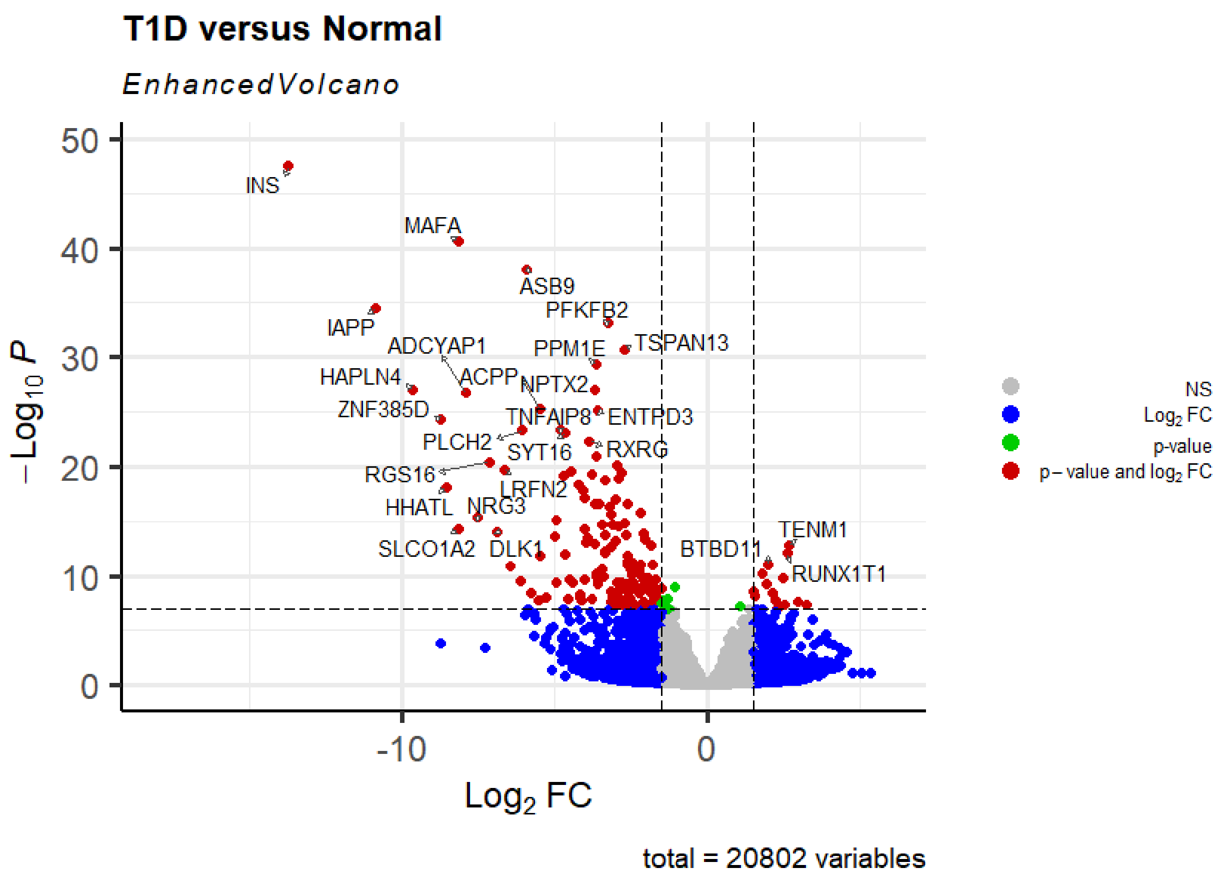

**Figure S5.** A dimensional reduction heatmap plot of the first 18 principal components (PCs) in the integrated dataset of Day1, Day29, $T_1D$, and $T_2D$.

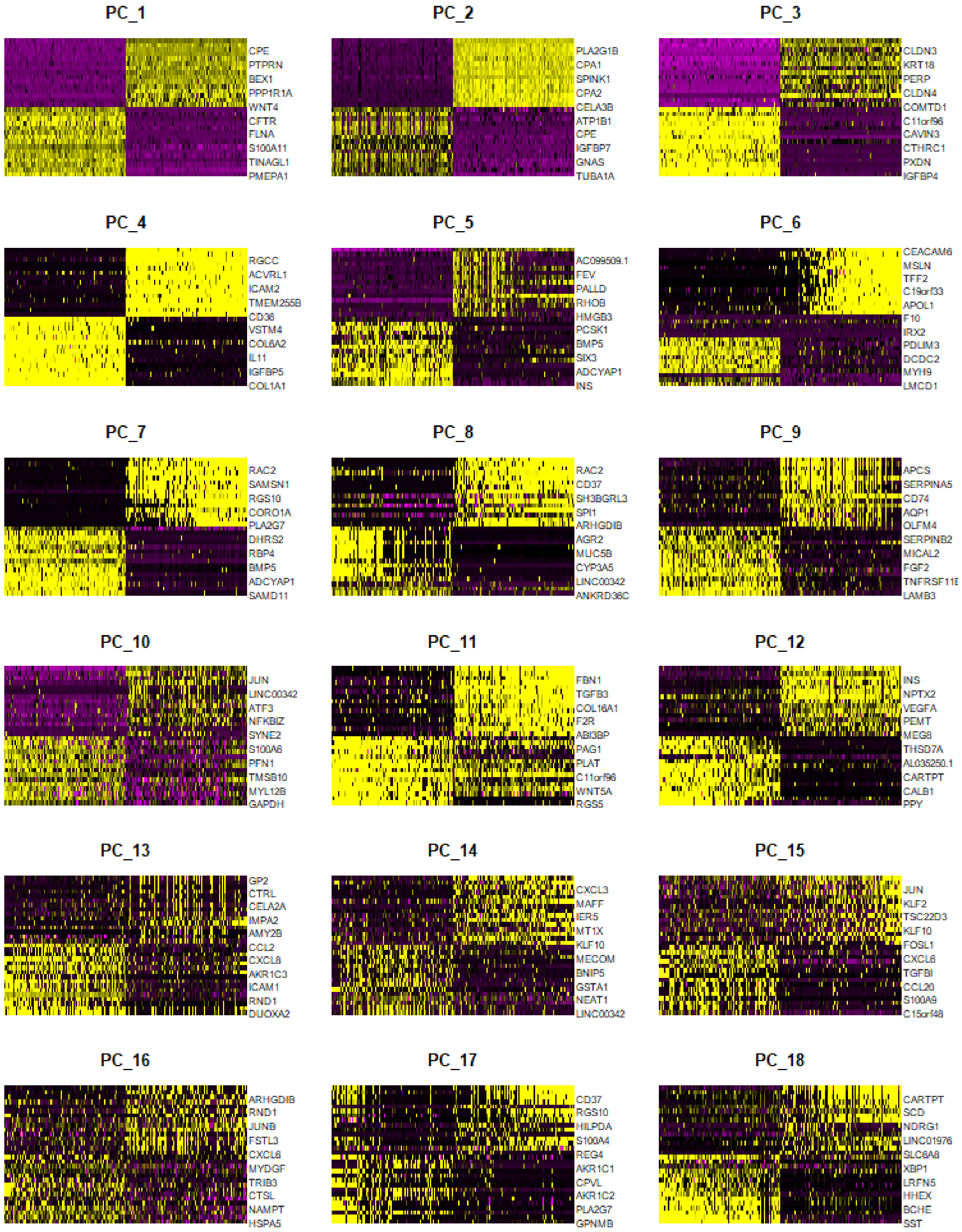

**Figure S6.** CIRCOS plots of summarized scRNA-seq data. **A.** A plot of clusters by cells counts in all samples. **B.** A plot of clusters by cells percentages in all samples

**A.** Numbers labeled on the left side of the plot represent the number of predicted cell clusters

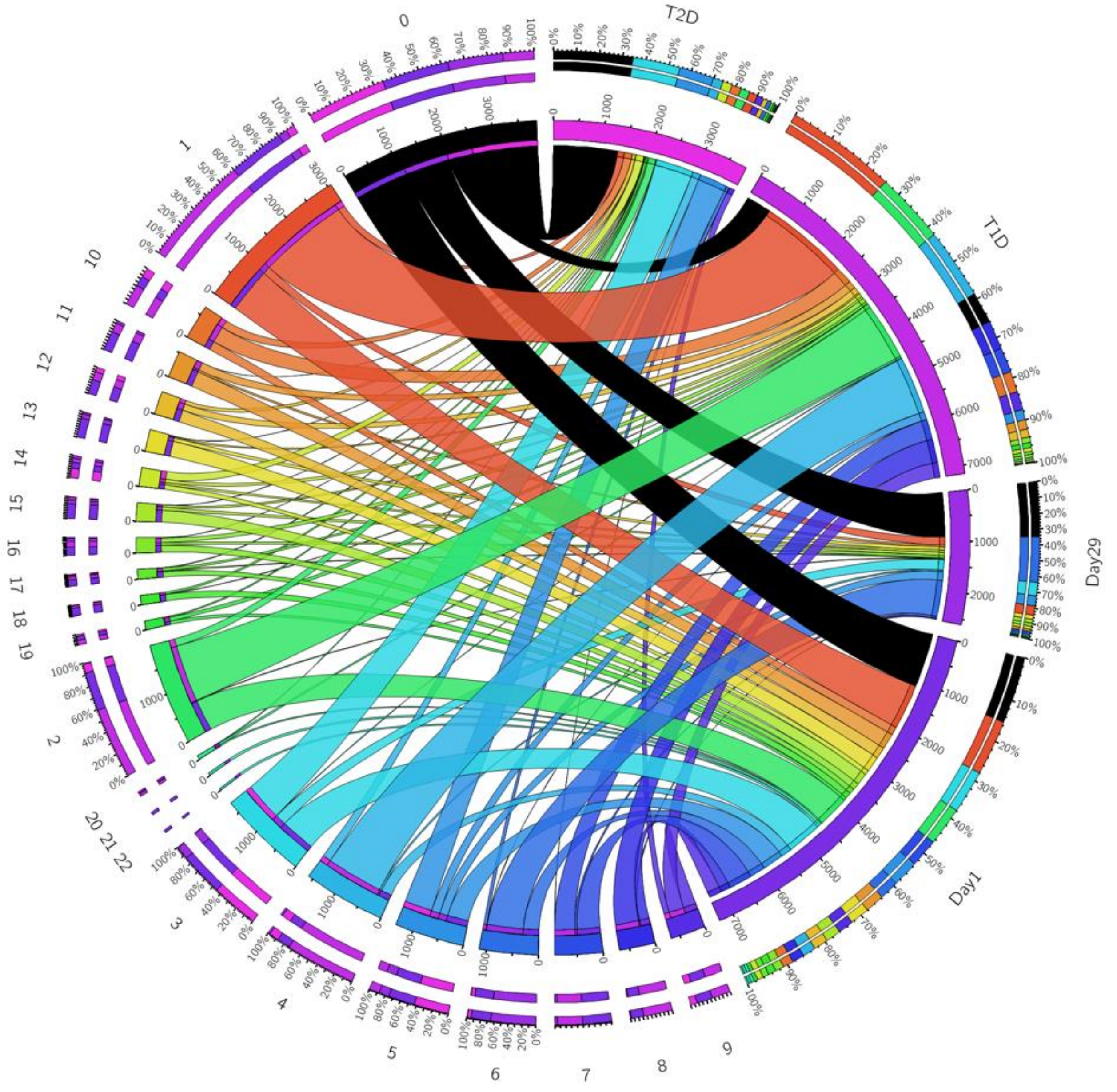

**B.** Numbers labeled on the left side of the plot represent the number of predicted cell clusters

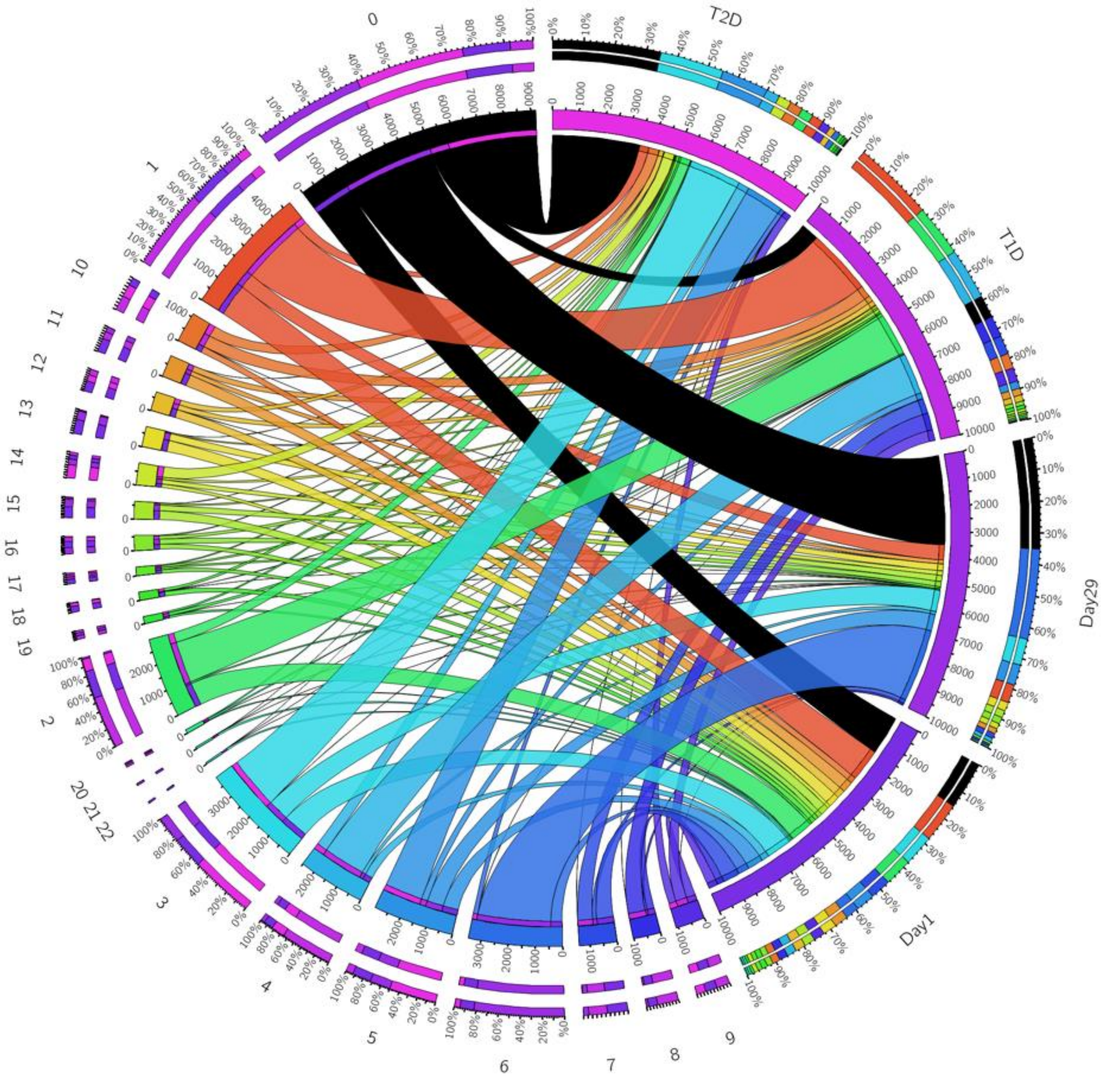

**Figure S7.** Ridge plots of expression ranges of *INS, GCG, MAFA*, and *MAFB* genes in each individual sample. **A.** The sample of Day1; **B.** The sample of Day29; **C.** Individual samples of T$_1$D; **D.** Individual samples of T$_2$D.

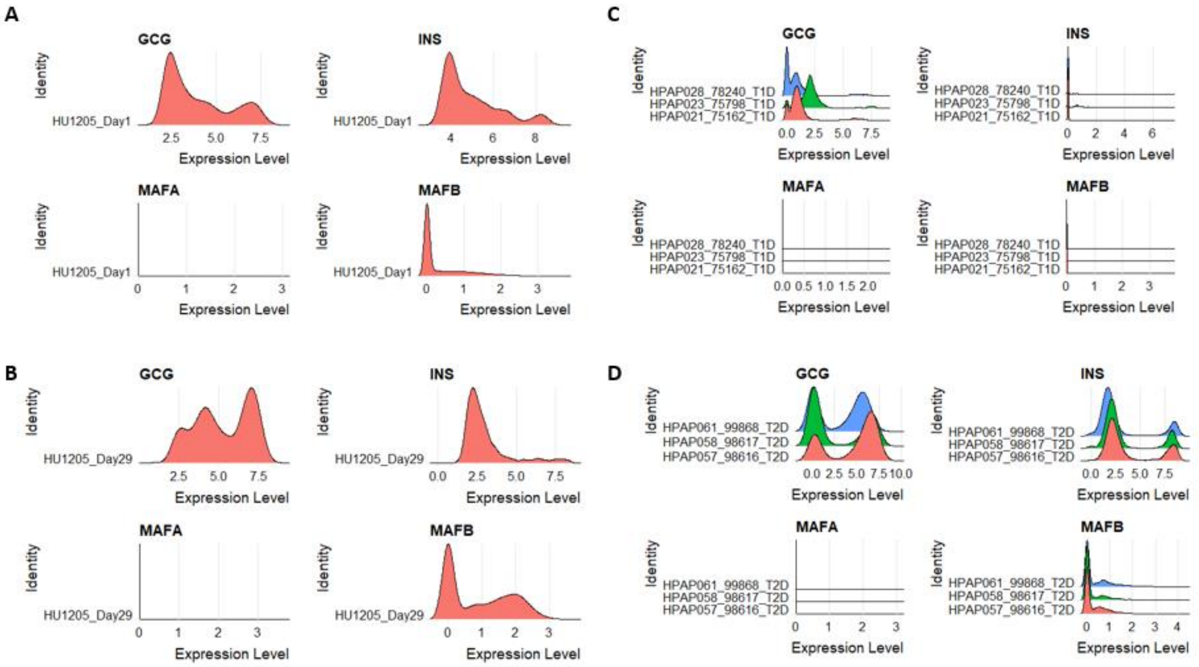

**Figure S8. DEG analysis of beta cells in cluster-3 and 6. A.** Venn diagram plot of too-many-cells analyzed DEGs between N19 (cluster-3) and N13 (cluster-6); **B.** GO enrichment from genes up-regulated in N19 vs N13 by too-many-cells; **C.** GO enrichment of genes down-regulated in N19 vs N13 by too-many-cells; **D.** Venn diagram plot of Seurat analyzed DEGs between N19 (cluster-3) and N13 (cluster-6) in Day1 or Day29 samples. **E.** Pathway analysis of down-regulated genes: cluster 6 vs cluster3 in Day1 sample by Seurat; **F.** Pathway analysis of up-regulated genes: cluster 6 vs cluster3 in Day1 sample by Seurat; **G.** Pathway analysis of down-regulated genes: cluster 6 vs cluster3 in Day29 sample by Seurat; **H.** Pathway analysis of up-regulated genes: cluster 6 vs cluster3 in Day29 sample by Seurat

**A.**

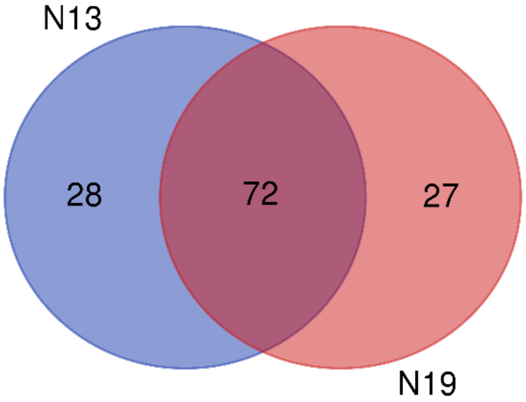


| Names | total | elements |
|---|---|---|
| N13 N19 | 72 | PGD LINC02593 AL031848.2 PIK3CD-AS2 AURKAIP1 RPL22 ENO1 CTNNBIP1 MRPL20 THAP3 AL390719.1 UBIAD1 MRPL20-AS1 LZIC FBXO2 SLC25A33 VAMP3 CENPS UBE4B SMIM1 CAMTA1 SRM CCNL2 FNDC10 TMEM240 CEP104 MAD2L2 LINC01128 PRDM2 CLSTN1 TARDBP TPRG1L AL645728.2 WRAP73 C1orf158 PEX10 B3GALT6 SDF4 FBXO44 DFFA PER3 PUSL1 PHF13 NPHP4 ICMT AL691432.2 DISP3 C1orf174 PEX14 SLC45A1 MIB2 UBE2J2 KCNAB2 SSU72 GNB1 KIAA2013 PARK7 RER1 AL109917.1 ENO1-AS1 ZBTB48 RERE LRRC47 NOC2L FAAP20 C1orf127 CDK11B ACOT7 ANKRD65 ATAD3C NMNAT1 SAMD11 |
| N13 | 28 | SLC35E2B KAZN INTS11 DVL1 NADK TNFRSF18 DFFB KIF1B CDK11A ATAD3A MIIP MTHFR FBXO6 NOL9 SKI FO704657.1 DNAJC11 EFHD2 VWA1 CASP9 LINC00115 MFN2 LINC01409 ISG15 MORN1 DNAJC16 AL645728.1 ARHGEF16 |

| | | |
|---|---|---|
| N19 | 27 | MUL1 FLJ37453 ACAP3 PINK1 UBXN10-AS1 TP73-AS1 OTUD3 UBXN10 PLEKHM2 UBR4 MICOS10 FAM131C UQCRHL CPLANE2 CAPZB SLC25A34 SPEN EMC1-AS1 FBXO42 SLC66A1 CAMK2N1 SZRD1 ZBTB17 AKR7A2 SDHB DDI2 EMC1 |

### B. N19 vs N13 ( Gene up-too-many-cells)

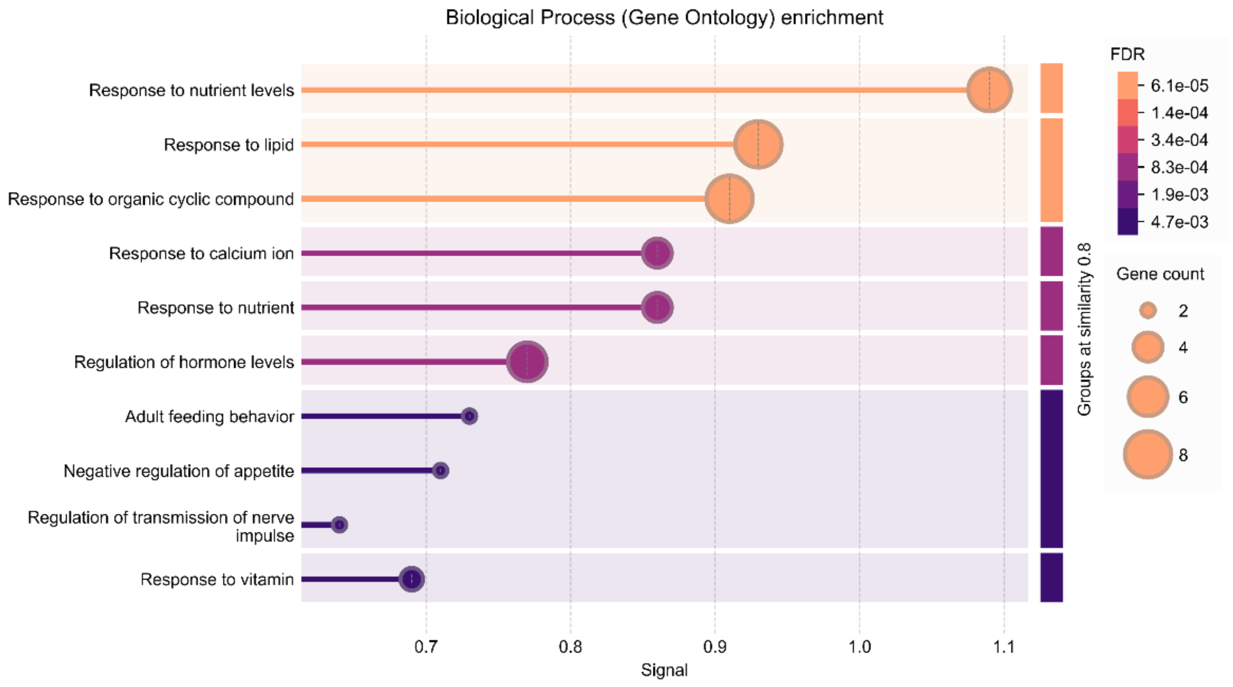


### C. N19 vs N13 ( Gene down)

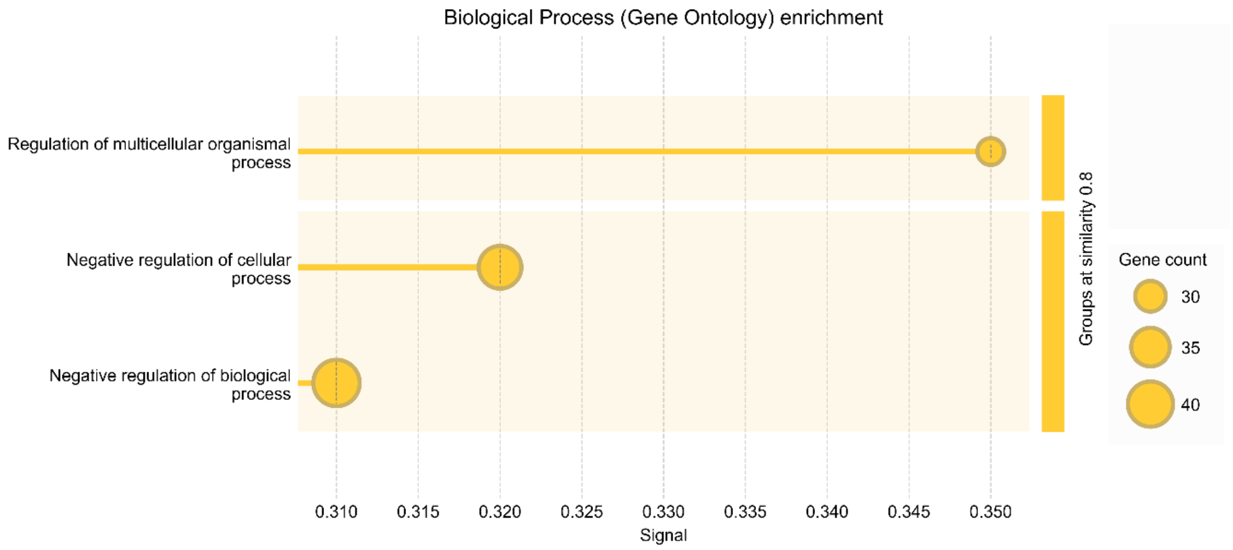

**D.** Venn diagram plot of DEGs of beta cells between cluster-6 and 3 by Seurat.

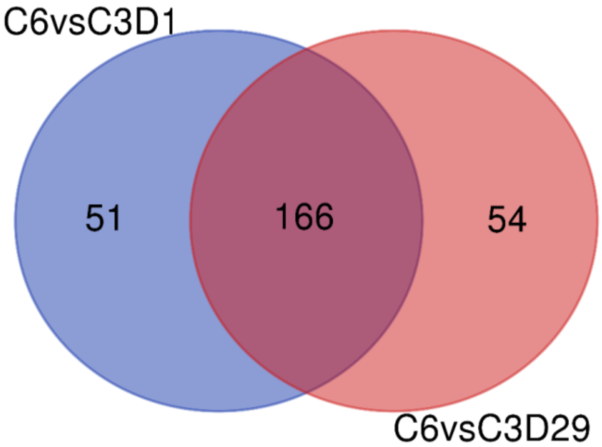


| Names | total | elements |
|---|---|---|
| C6_vs_C3_Day1 C6_vs_C3_Day29 | 166 | MT1G CD44 KCNK16 SCG3 PNLIP FXYD2 SPINK1 CPA1 MAP1B ARG2 MANF ERO1B KCNQ1OT1 NPTX2 PPY MAP1LC3A CITED2 TMOD1 INS FAM162A TRIB3 LOXL4 CLGN SOD2 GSN SYNE2 OSTC MEG3 PEMT SELENOW CDKN1A PCSK1 DNAJB1 TUBA1A FAP SORL1 REG1B DDIT4 LUCAT1 TCEAL2 LMO2 STX1A INSM1 RBP4 ADCYAP1 PCP4 PAPSS2 SCGN MPP1 CTRC SLC30A8 NUCB2 GADD45A IAPP PPP1R1A KRT10 PLA2G1B ELAVL4 ENO2 UCHL1 SCG2 TMEM236 GNAS CFC1 TSPAN13 GNG4 CLDN3 PAK3 FOSB CLPS DHRS7 SHISAL2B CPE GAD2 VGF ANKRD36C NR4A1 AQP3 ATP1B1 PAPPA2 WNT4 PDX1 SAMD11 CACNA1A SYT13 NEUROD1 DDIT3 TXN SDF2L1 HSPA5 LMO1 DLK1 COMTD1 NKX6-1 SNHG7 SLC6A6 C12orf75 HADH PRUNE2 C1QL1 GAPDH CPB1 VEGFA DNAJC12 PARVB ABRACL OTULINL CDKN2A GRAMD4 ABCC8 QPCT STMN2 MT1X CELA3A APLP2 HSD17B14 CTRB1 PRDX1 CTSL SEC11C ELMO1 SCGB2A1 BMP5 MYDGF DNAJB9 PRDX4 PRSS23 CHPF SIPA1L2 TFF3 HSP90B1 DEPP1 BAG1 PDK4 NEAT1 RASD1 HSPA8 BEX1 MAFA LAPTM4B CEL PTS MAL2 PKIB TMEM37 ANXA5 BTG3 SIX3 RETREG1 DUSP26 REG1A PERP XBP1 CASR COX7A1 PCSK1N HOPX FTL MAFB CADM1 KLF6 RNF213 DHRS2 OLFM2 C4orf48 EEF1A2 |
| C6_vs_C3_Day1 | 51 | TKT GAS5 MT1F JUND SH3BGRL3 MKNK2 MYL12B ARL6IP1 TSC22D1 GSTO1 HERPUD1 SCD SERPINE2 UGDH IFI6 CNIH2 SDCBP ID4 NDUFA4 B2M GPX3 TMEM45A TMEM176A PFN1 GAMT TIMP2 CD200 HEXB NPY QDPR SST TPPP3 SELENOM PLCXD3 JUNB CTSZ CYB5A ID1 NKX2-2 GEM TSPAN7 RBP1 SMOC1 TNFRSF12A CAMK2G TMEM176B ANXA11 PDE10A MPC1 SLC7A2 HSPH1 |

| | | |
|---|---|---|
| C6_vs_C3_Day29 | 54 | TM7SF2 KCNMA1 BTG2 CEP126 CHGB COL4A1 PAM COL4A2 COL6A2 EDIL3 IFITM3 APCDD1L-DT IRF1 IL32 KLF2 VIM TUBB2B GCG INHBA ADM PALLD OLMALINC COL1A1 ATF3 YBX3 ZFP36L1 LGALS1 CALD1 RAB3B C1orf127 C11orf96 NFKBIA CRELD2 MEIS2 IGFBP4 COL18A1 MYH9 ELF3 F10 ACSL1 ABHD3 IRX2 IGFBP7 FLNA MACC1 SYT4 PTPRN SAT1 ATP2A3 GNG11 ODC1 EGFL7 WLS CD14 |

**E.** Down-regulated genes: cluster 6 vs cluster3 in Day1 sample by Seurat

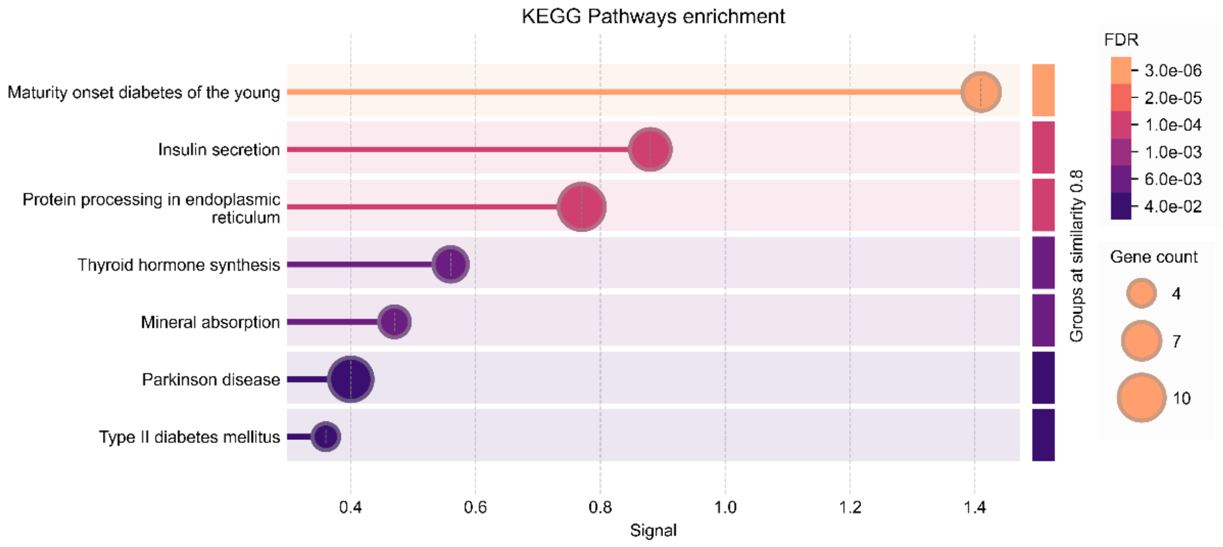

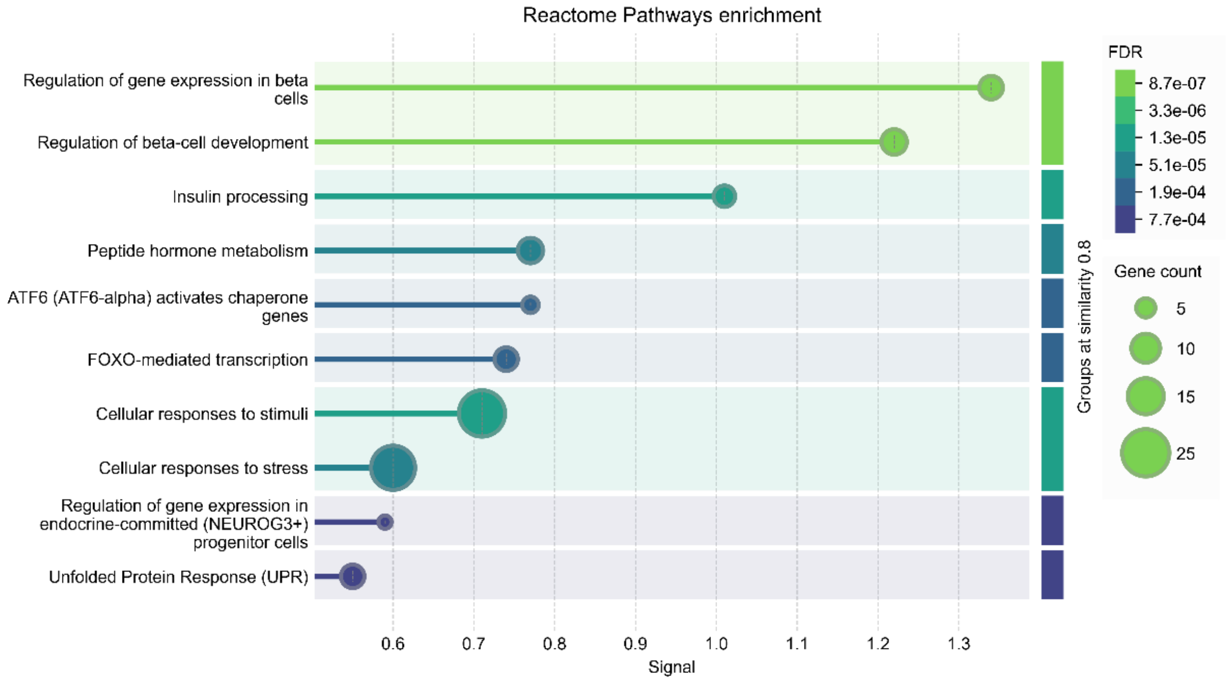

Reactome Pathways enrichment
Regulation of gene expression in beta cells
Regulation of beta-cell development
Insulin processing
Peptide hormone metabolism
ATF6 (ATF6-alpha) activates chaperone genes
FOXO-mediated transcription
Cellular responses to stimuli
Cellular responses to stress
Regulation of gene expression in endocrine-committed (NEUROG3+) progenitor cells
Unfolded Protein Response (UPR)
0.6
0.7
0.8
0.9
1.0
1.1
1.2
1.3
Signal
Groups at similarity 0.8
FDR
8.7e-07
3.3e-06
1.3e-05
5.1e-05
1.9e-04
7.7e-04
Gene count
5
10
15
25


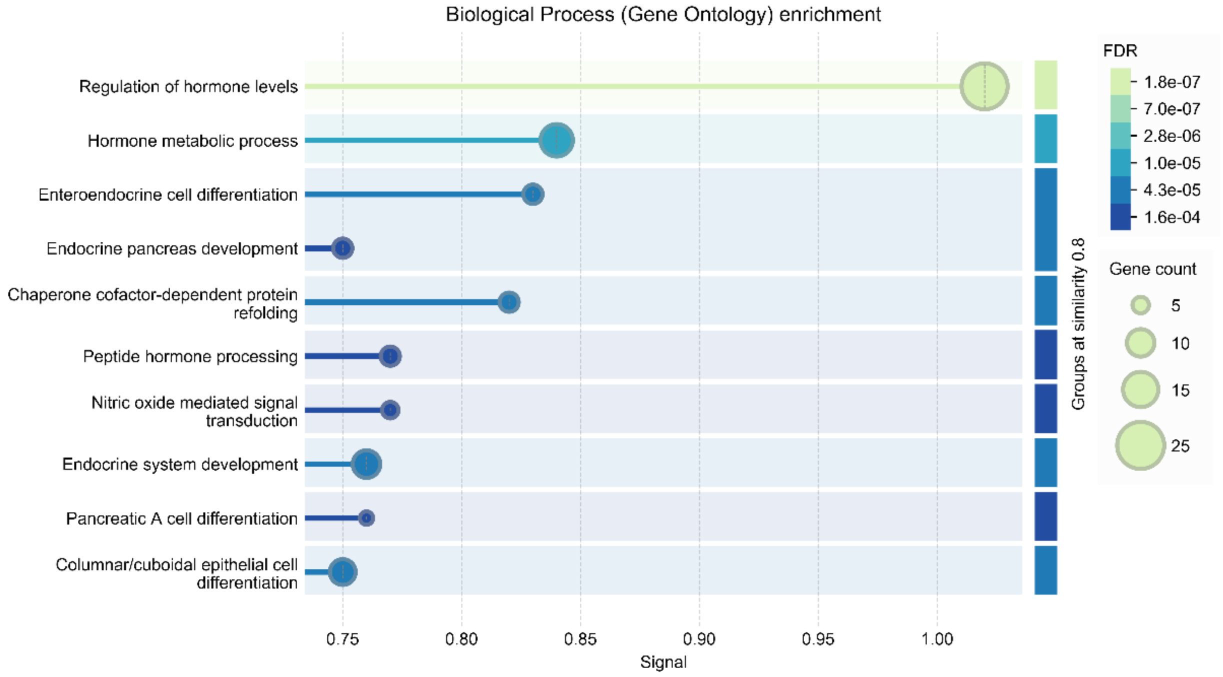

Biological Process (Gene Ontology) enrichment
Regulation of hormone levels
Hormone metabolic process
Enteroendocrine cell differentiation
Endocrine pancreas development
Chaperone cofactor-dependent protein refolding
Peptide hormone processing
Nitric oxide mediated signal transduction
Endocrine system development
Pancreatic A cell differentiation
Columnar/cuboidal epithelial cell differentiation
0.75
0.80
0.85
0.90
0.95
1.00
Signal
Groups at similarity 0.8
FDR
1.8e-07
7.0e-07
2.8e-06
1.0e-05
4.3e-05
1.6e-04
Gene count
5
10
15
25

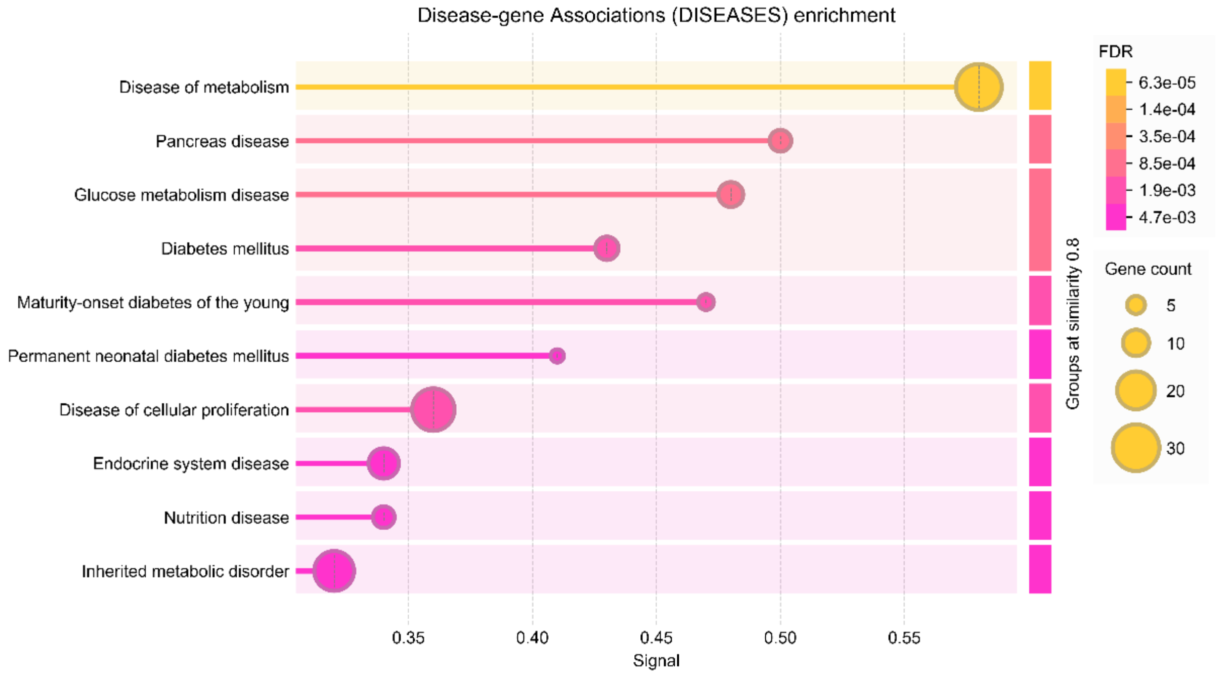


**F.** Up-regulated genes: cluster 6 vs cluster3 in Day1sample by Seurat

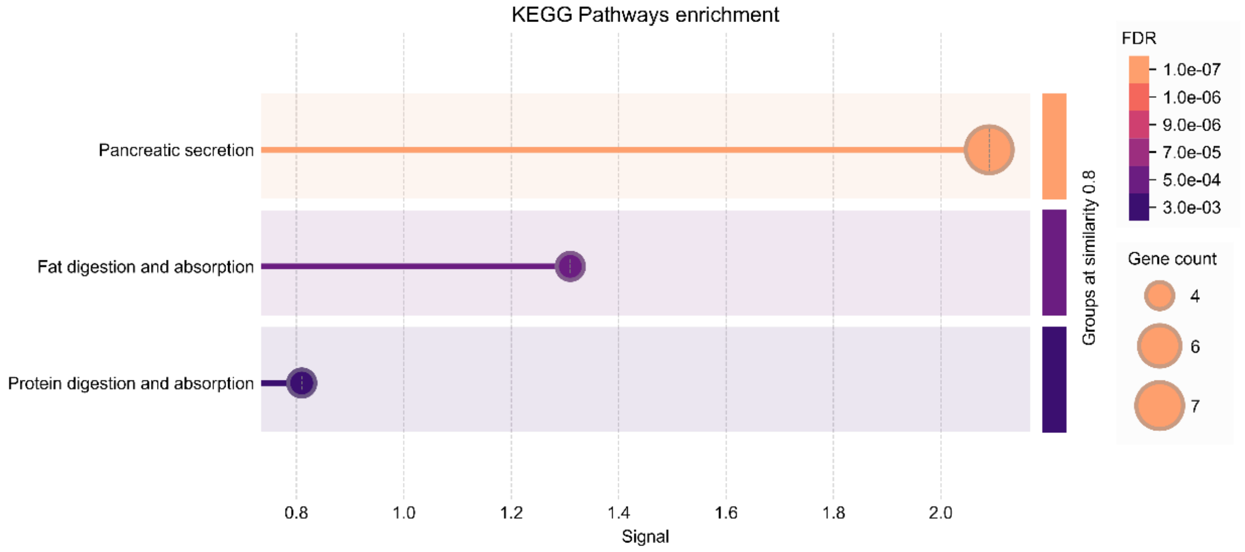

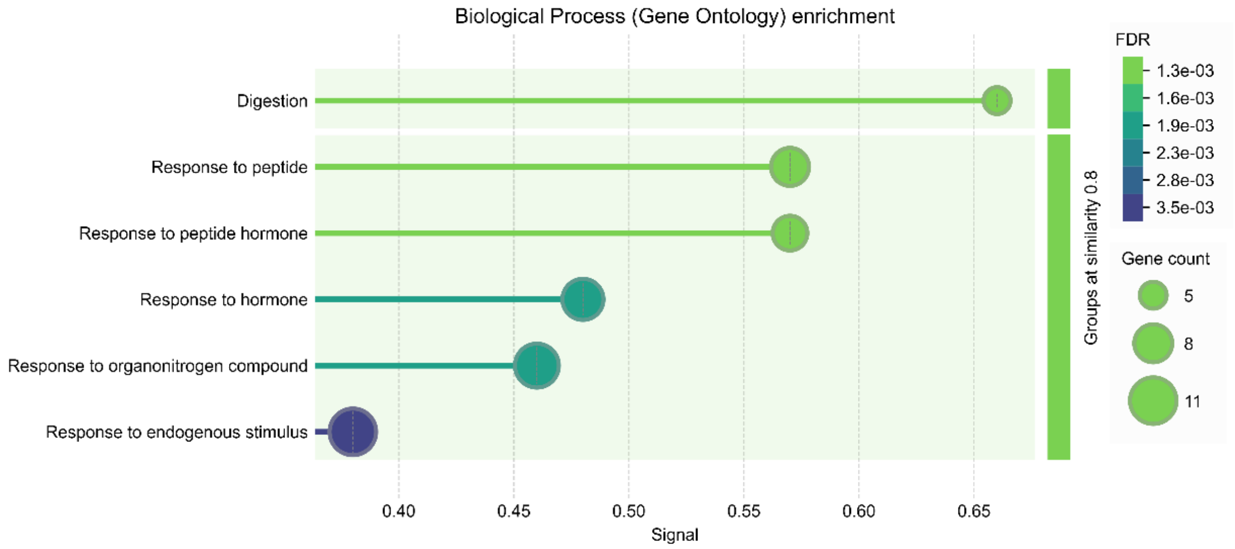


**G.** Down-regulated genes: cluster 6 vs cluster3 in Day29 sample by Seurat

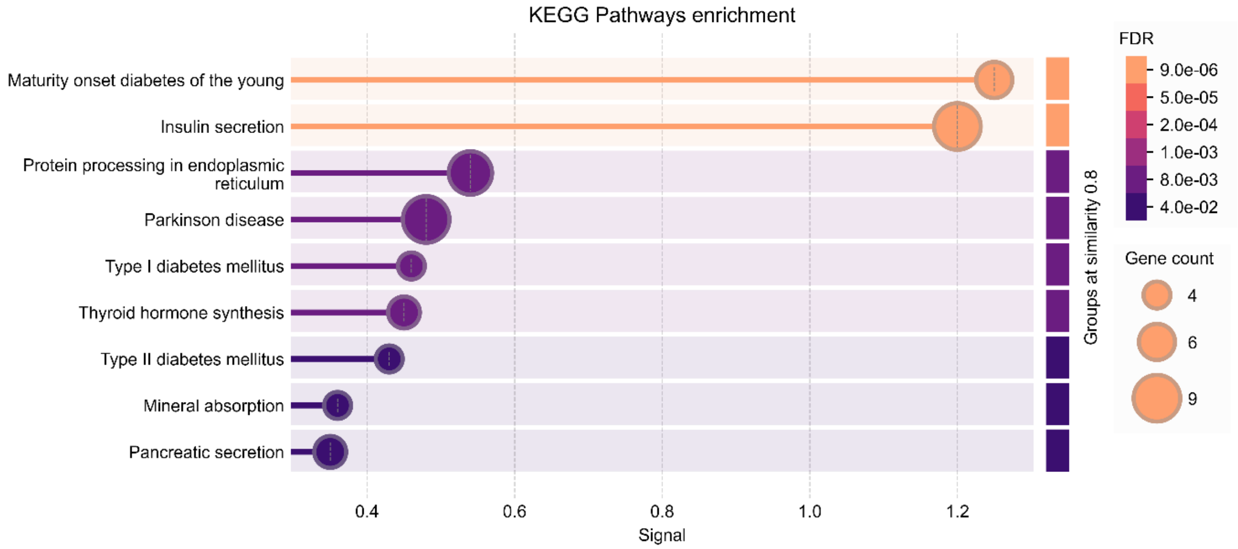

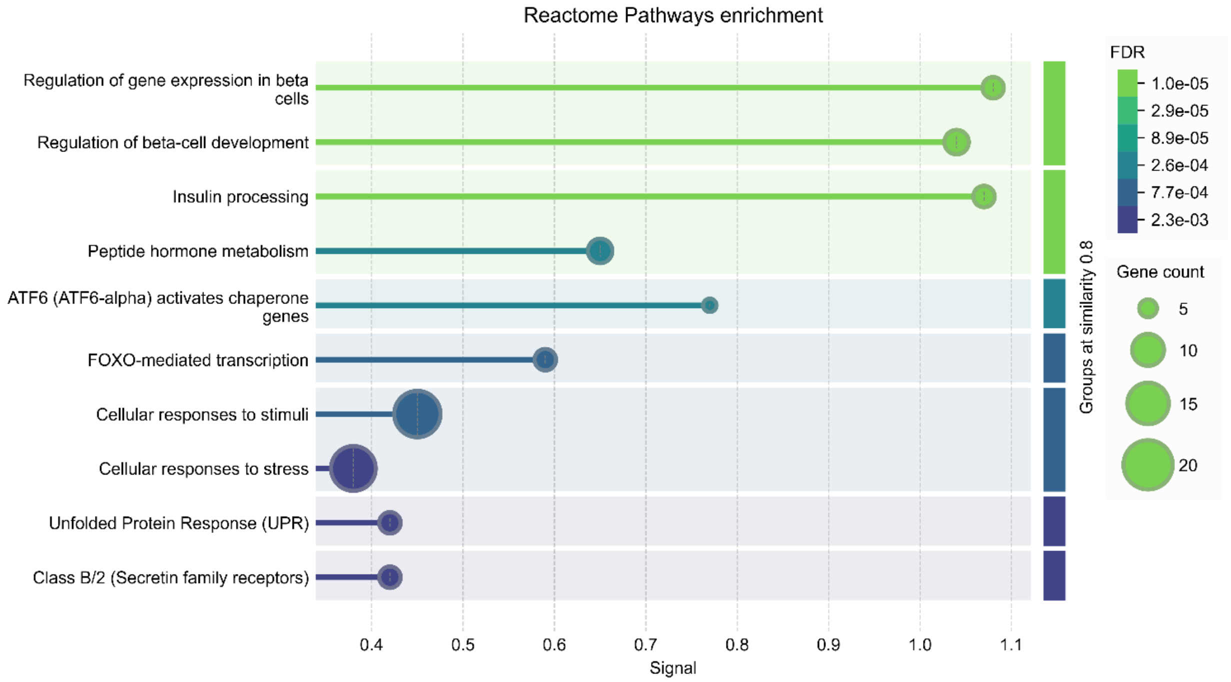

Reactome Pathways enrichment
Regulation of gene expression in beta cells
Regulation of beta-cell development
Insulin processing
Peptide hormone metabolism
ATF6 (ATF6-alpha) activates chaperone genes
FOXO-mediated transcription
Cellular responses to stimuli
Cellular responses to stress
Unfolded Protein Response (UPR)
Class B/2 (Secretin family receptors)
0.4
0.5
0.6
0.7
0.8
0.9
1.0
1.1
Signal
Groups at similarity 0.8
FDR
1.0e-05
2.9e-05
8.9e-05
2.6e-04
7.7e-04
2.3e-03
Gene count
5
10
15
20


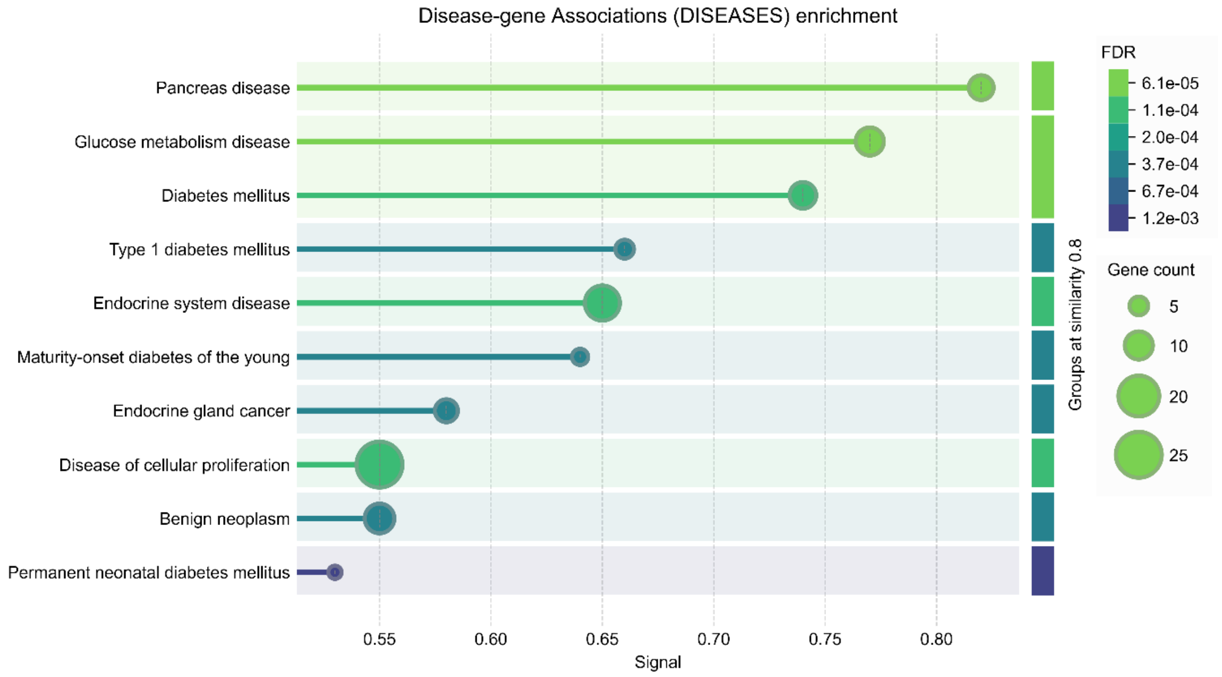

Disease-gene Associations (DISEASES) enrichment
Pancreas disease
Glucose metabolism disease
Diabetes mellitus
Type 1 diabetes mellitus
Endocrine system disease
Maturity-onset diabetes of the young
Endocrine gland cancer
Disease of cellular proliferation
Benign neoplasm
Permanent neonatal diabetes mellitus
0.55
0.60
0.65
0.70
0.75
0.80
Signal
Groups at similarity 0.8
FDR
6.1e-05
1.1e-04
2.0e-04
3.7e-04
6.7e-04
1.2e-03
Gene count
5
10
20
25

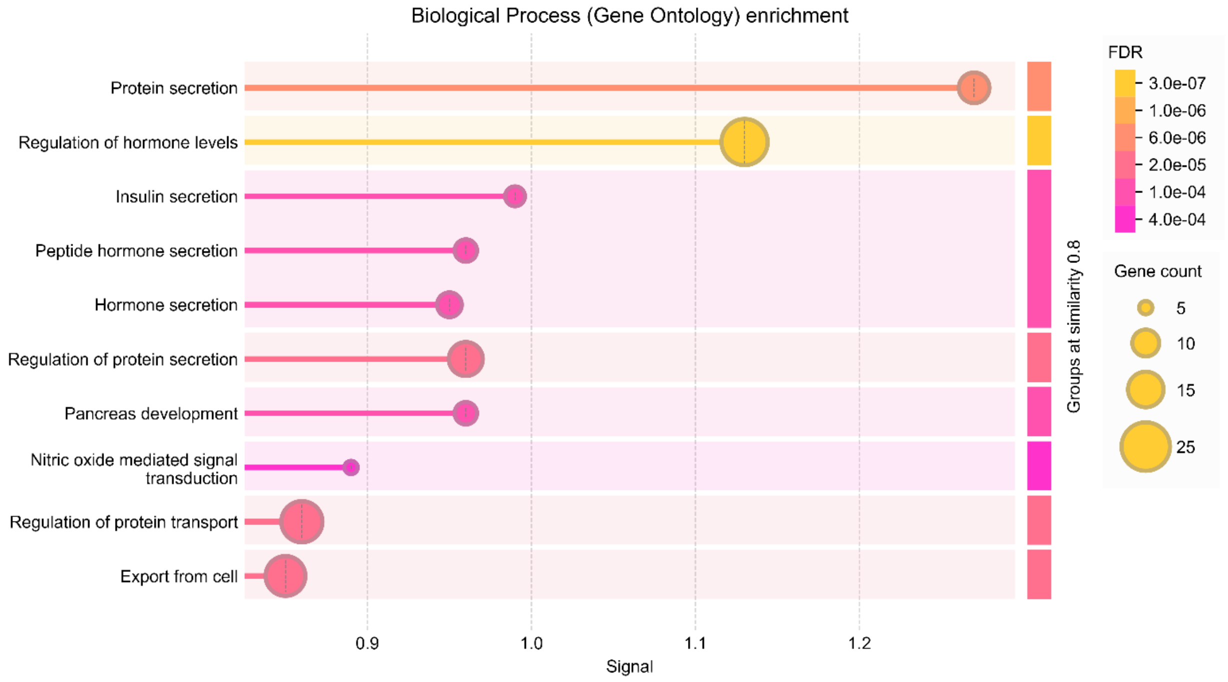


**H.** Up-regulated genes: cluster-6 vs cluster3 in Day29 sample by Seurat

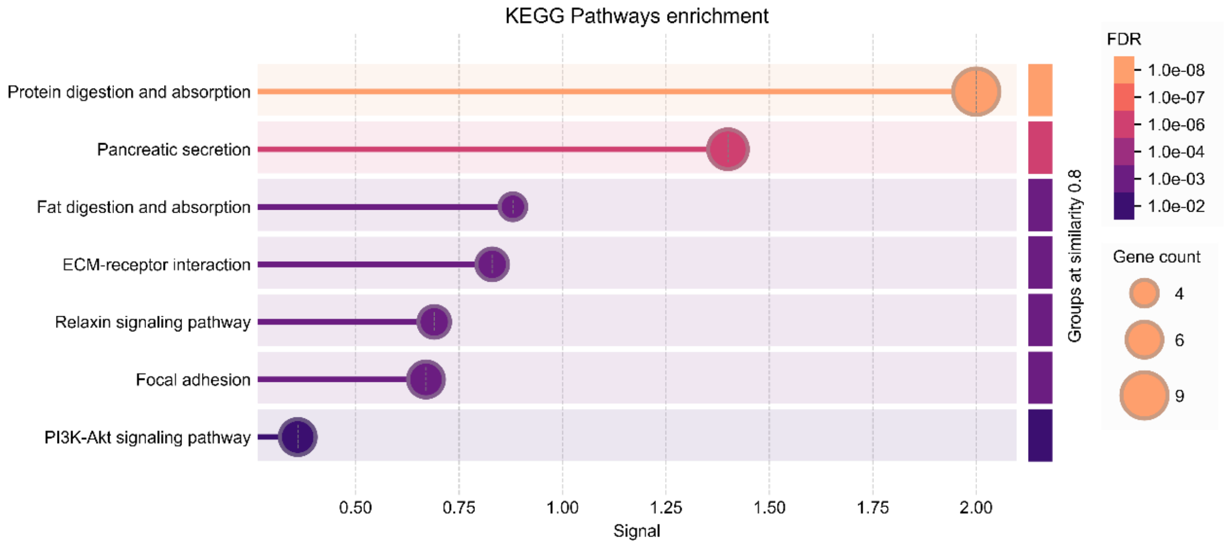

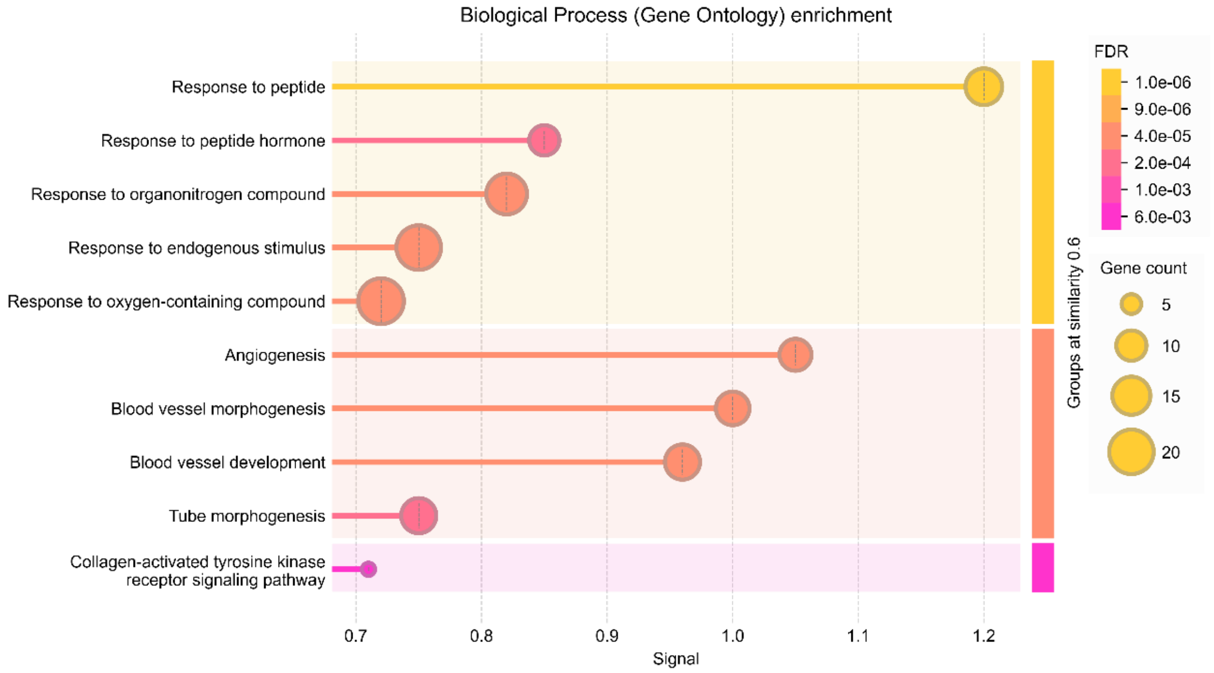
Biological Process (Gene Ontology) enrichment
Response to peptide
Response to peptide hormone
Response to organonitrogen compound
Response to endogenous stimulus
Response to oxygen-containing compound
Angiogenesis
Blood vessel morphogenesis
Blood vessel development
Tube morphogenesis
Collagen-activated tyrosine kinase receptor signaling pathway
0.7
0.8
0.9
1.0
1.1
1.2
Signal
Groups at similarity 0.6
FDR
1.0e-06
9.0e-06
4.0e-05
2.0e-04
1.0e-03
6.0e-03
Gene count
5
10
15
20

**Figure S9.** CellChat predicted number of interactions and strength of interactions between different cell groups**. A.** The sample of Day1; **B.** The sample of Day29; **C.** The sample of $T_1D$; **D.** The sample of $T_2D$.

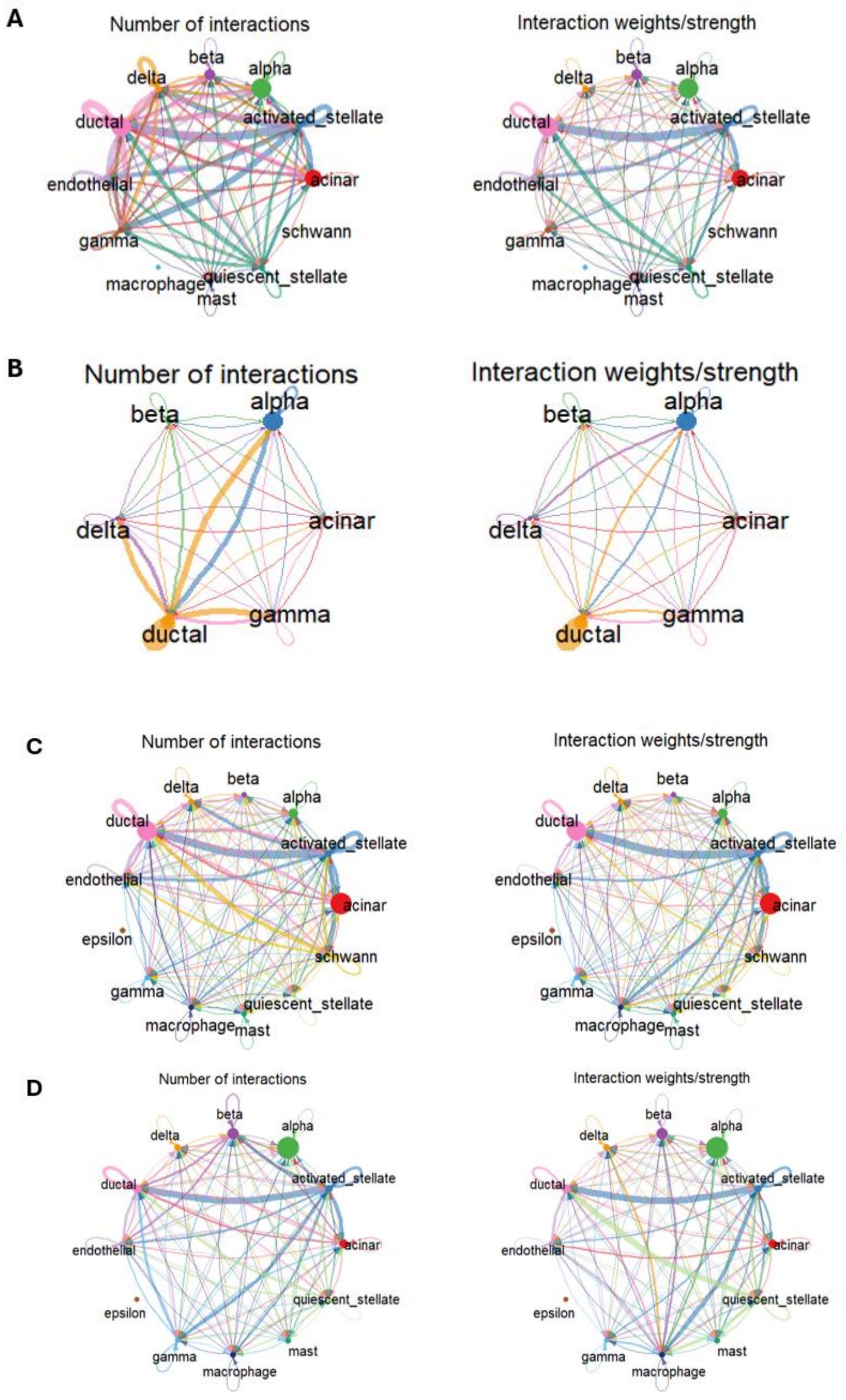
